# Prevalence of mutualism in a simple model of microbial co-evolution

**DOI:** 10.1101/2021.09.03.458878

**Authors:** Luciano Stucchi, Javier Galeano, Juan Manuel Pastor, Jose María Iriondo, José A. Cuesta

## Abstract

Evolutionary transitions among ecological interactions are widely known, although their detailed dynamics remain absent for most population models. Adaptive dynamics has been used to illustrate how the parameters of population models might shift through evolution, but within an ecological regime. Here we use adaptive dynamics combined with a generalised logistic model of population dynamics to show that transitions of ecological interactions might appear as a consequence of evolution. To this purpose we introduce a two-microbial toy model in which population parameters are determined by a bookkeeping of resources taken from (and excreted to) the environment, as well as from the byproducts of the other species. Despite its simplicity, this model exhibits all sorts of ecological transitions, some of which resemble those found in nature. Overall, the model shows a clear trend toward the emergence of mutualism.

## Introduction

In 1966, Jeon and Lorch were conducting experiments with a population of *Amoeba proteus*. One of the strains got infected by X-bacteria, a Gram-negative, rod-shaped bacteria related to *Legionella* sp. and *Pseudomonas* sp. (for which the name *Candidatus Legionella jeonii* sp. nov. was later proposed (***Park et al., 2004***)). The few survivors to the infection retained the bacteria as parasites (***Jeon and Lorch, 1967***). Over the course of a few generations though, they became endosymbionts, providing amoebas heatshock protection (***Jeon, 1992, 1995***).

Arguably, this is one of the most spectacular cases of an evolutionary transition in an ecosystem from an antagonistic interaction to an obligate mutualism, mainly because it was witnessed in real time. However, other similar transitions are well documented in the literature from different sources. For instance, phylogenetic data collected from one of the best-known mutualistic systems—plants and pollinators—dating back 90 million years, allowed ***Machado et al. (2001***) to show that the wasp *Ceratosolen galili*, which belongs to a genus of active pollinating species, not only does not pollinate anymore but has become a parasitic species of figs.

Likewise, performing phylogenetic analysis on 15 species of aphids of the genus *Chaitophorus*, ***Shingleton and Stern (2002***) concluded that the relationship between ants and aphids may have changed at least five times during the history of their life, thus going through all shades between mutualism and antagonism. There is no consensus whatsoever on whether the ancestral mutualistic relationship between these species was facultative or obligatory, but it seems reasonable to think that, before these species developed any mutualistic relationship, ants predated on aphids because this behaviour is still observed in all ant species in the appropriate environmental conditions (***Sakata, 1994; Stadler and Dixon, 2005; Offenberg, 2001***).

The above-mentioned examples are only a few well-studied ecological systems that show transitions between different types of ecological interactions. In all of them, the evolutionary nature of these transitions has been established either through experiments or using phylogenetic analyses (***Sachs and Simms, 2006***), but their presence suggests that many more may have occurred in nature. This fact raises many questions: Are these evolutionary transitions a common phenomenon or a rarity? Why do ecological interactions move in one direction or another? Would it be possible to predict in which direction a type of interaction will move, if at all?

One way to explore the answers to some of these questions is to design mathematical models that include two different time scales: a short one that accounts for the usual population dynamics, and a long one that includes Darwinian evolution. The latter can be accomplished by applying adaptive dynamics to the parameters of the population model (***Dieckmann and Law, 1996; Dercole and Rinaldi, 2008; Doebeli, 2011***). However, the former is more problematic because standard population models deal with different types of interactions in very different ways.

When it comes to modelling competition and predation, Lotka-Volterra population equations are a very convenient choice (***Lotka, 1925; Volterra, 1926, 1928***). Despite their simplicity, these equations yield a rich set of biological predictions, to the point that they can be used as a prototype for more realistic models as well as a tool for interpreting complex observations. However, the Lotka-Volterra model meets serious difficulties to accommodate mutualism because positive interactions between species induce spurious feed-back loops that may drive populations out of control (***May, 1981***). The addition of Holling-type functional responses (***Wright, 1989***) is the usual way to control an unbounded growth of the populations, at the expense of rendering the model analytically intractable. Recently though, a new population model has been introduced in which both, species’ vegetative growth and interspecific interactions, are limited via logistic terms (***García-Algarra et al., 2014; Stucchi et al., 2020***). The resulting equations are amenable to analytic treatment and, at the same time, provide sensible results whether the interactions are of antagonistic or mutualistic nature. These two nice features make this model particularly suitable to study the evolution of ecological interactions.

Variation in ecological interactions may take place as a result of multiple factors (***Thompson, 1988***). For instance, the outcomes of ecological interactions may depend on the age and/or life cycle stage of the individuals of the interacting species. Similarly, phenotypic differences between the individuals of each of the interacting species, may alter the impact of one species on the other. These differences can result from genotypic diversity in the interacting populations and/or phenotypic plasticity. In order to focus on the general patterns derived from population dynamics and evolutionary adaptation, we have chosen a basic microbial system in which there are no differences among the individuals of each of the interacting species as a result of age or life cycle stage. Furthermore, there is one single genotype per interacting species and the environmental conditions are fixed. Therefore, all individuals of each interacting species have the same phenotype. Also for simplicity we will assume that mutations are so rare that they either get quickly fixed in the whole population or disappear before a new one occurs. In this regime, a suitable theoretical tool to study evolution is adaptive dynamics (***Dieckmann and Law, 1996; Dercole and Rinaldi, 2008; Doebeli, 2011***).

Accordingly, our toy model consists of two microbial species that interact via the resources they consume and excrete. They compete for resources from the environment—when both of them use the same resource—but they can also cross-feed from resources excreted by the other species. This interplay between resource consumption and excretion will provide specific functional forms for the parameters of ***Stucchi et al***.’s population equations, automatically yielding natural trade-offs between these parameters (***Tilman, 2004***). Adaptive dynamics will then introduce the Darwinian mechanism for the evolution of the parameters of the model.

Simplistic as it may be, this model provides sufficient complexity and flexibility as to show evolutionary transitions of all sorts. It reveals a global trend toward the appearance of mutualistic interactions from initially competitive or antagonistic scenarios, and exhibits evolutionary pathways akin to some documented in the literature.

In what follows we provide a detailed account of the model for two species, starting with a brief description of ***Stucchi et al***.’s ecological model and further connecting its phenomenological parameters with the microscopic interactions between the species. The equations of the adaptive dynamic for this model are developed in the ***Appendix 1***. We end by providing and analysing the results of extensive numerical simulations.

## Results

### A toy model for two interacting microbial species

In this section we will introduce what might be the simplest possible model capable of exhibiting evolutionary transitions between different ecological regimes. We will consider two microbial species whose populations evolve in time according to the *generalised logistic model* (***Stucchi et al., 2020***), briefly described in ***Box 1***. Specialising this model for just two species, and ignoring intraspecific cooperation or direct competition (i.e., *b*_*ii*_ = 0), the two equations that describe this minimal ecological community are

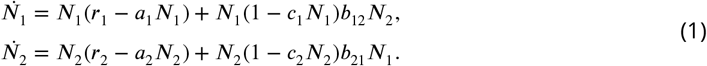

This system of differential equations is able to describe every kind of ecological interaction. ***Table 1*** shows the kind of relationships of the species with the environment (through the signs of the vegetative growth rates *r*_*i*_) and between them (through the signs of the interaction constants *b*_*ij*_) which correspond to the most usual ecological scenarios. Depending on the signs of *b*_*ij*_ the interaction between both species can be *mutualistic* (both positive), *antagonistic* (one positive and one negative, also called *predation* or *parasitism*, depending on the context), or *competitive* (both negative). On the other hand, the relation with the environment determines whether mutualism is *facultative* (*r*_*i*_ > 0) or *obligate* (*r*_*i*_ < 0). The case of *predation/parasitism* (*b*_*ij*_ < 0, *b*_*ji*_ > 0, i.e. species *j* benefits at the expense of species *i*) requires the prey or parasitised species to obtain resources from the environment (*r*_*j*_ > 0) for the system to be sustainable. When *b*_*ij*_ ≈ 0, this sort of interaction is usually referred to as *commensalism*. Finally, *competition* (*b*_*ij*_ < 0, *b*_*ji*_ < 0) occurs when both species are a hindrance to each other (which of course needs *r*_*i*_ > 0 and *r*_*j*_ > 0 for the system to be sustainable).

**Table 1.** Signs of *r*_*i*_ and *b*_*ij*_ that describe typical ecological interactions.

| $r_i$ | $r_j$ | $b_{ij}$ | $b_{ji}$ | type of interaction | abbreviation |
| --- | --- | --- | --- | --- | --- |
| + | + | + | + | facultative-facultative mutualism | FFM |
| + | - | + | + | obligate-facultative mutualism | OFM |
| - | - | + | + | obligate-obligate mutualism | OOM |
| + | + | - | - | competition | CMP |
| + | + | - | + | facultative predator-prey / parasitism | FPP |
| + | - | - | + | obligate predator-prey / parasitism | OPP |
| + | any | $\sim 0$ | + | commensalism | COM |

Parameters *r*_*i*_ and *b*_*ij*_ are purely phenomenological. However, in a real system, they depend on the specific ways in which the species relate with the environment and with each other. Because of this, the parameters are not independent. However, to specify how they are connected, we need to delve into the details of the system we want to model. This is a necessary step to construct an evolutionary model of an ecosystem (***Dieckmann and Law, 1996***).

Let us assume that the two microbial species described by ***Equation 1*** struggle to survive in a fixed environment (described by the parameters *a*_*i*_ and *c*_*i*_), with a given number *K* of available resources. Each species consumes *n*_*i*_ of these *K* resources, *q* of which are common to both species. As a consequence of internal metabolic reactions, species *i* produces *m*_*i*_ byproducts, *m*_*ji*_ of which are useful to the other species *j* and *ℓ*_*i*_ = *m*_*i*_ − *m*_*ji*_ are not. Even though *m*_*i*_ depends on the details of the metabolism of species *i*, for the sake of simplicity we can just assume that it is proportional to the number of consumed resources, i.e.

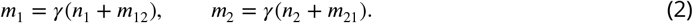

A fraction 0 < *α*_*j*_ < *γ* of species *i*’s metabolic byproducts will be used by species *j*; hence,

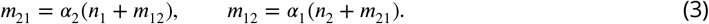

Coefficients *α*_*i*_ can evolve over time. ***Equation 3*** is a linear system whose solution is

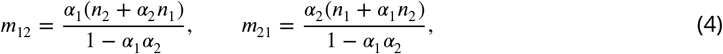

and substituting in ***Equation 2***,

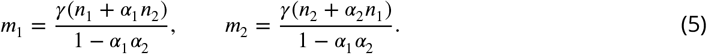

These equations express the number of metabolic residues as a function of the number of resources.

Finally, we need to connect the demographic parameters with this flux of resources and byproducts. In principle, microbes fare better the more resources they have, whereas metabolic byproducts are costly. Thus, we posit that the growth rate increases proportional to the amount of consumed resources and decreases with the amount of metabolic byproducts, i.e.

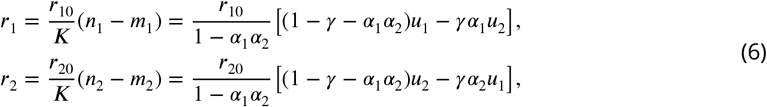

denoting *u*_*i*_ ≡ *n*_*i*_/*K*. Notice that the factor *K* is introduced for convenience, so as to express everything in terms of the re-scaled variables 0 ⩽ *u*_*i*_ ⩽ 1.

Likewise, the interaction coefficients *b*_*ij*_ increase with *m*_*ij*_, the number of byproducts of species *j* that are useful to species *i*, and decrease with the competition for the shared resources *q*. Hence,

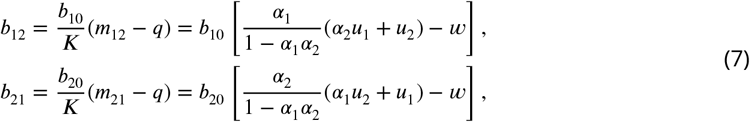

where we have denoted *w* ≡ *q*/*K* (0 ⩽ *w* ⩽ min(*u*_1_, *u*_2_)).

Parameters *r*_*i*0_ and *b*_*i*0_ are simple dimensional constants that also set the time scale of the differential equations.

Thus, after these simplifying assumptions, we end up with a model in which the demographic parameters are expressed in terms of the number of consumed resources, *n*_1_, *n*_2_, the resource sharing *q*, and the cross-feeding efficiencies *α*_1_, *α*_2_. A sketch of the model, where all this interactions are summarised, is shown in ***Figure 1***.

**Figure 1.**
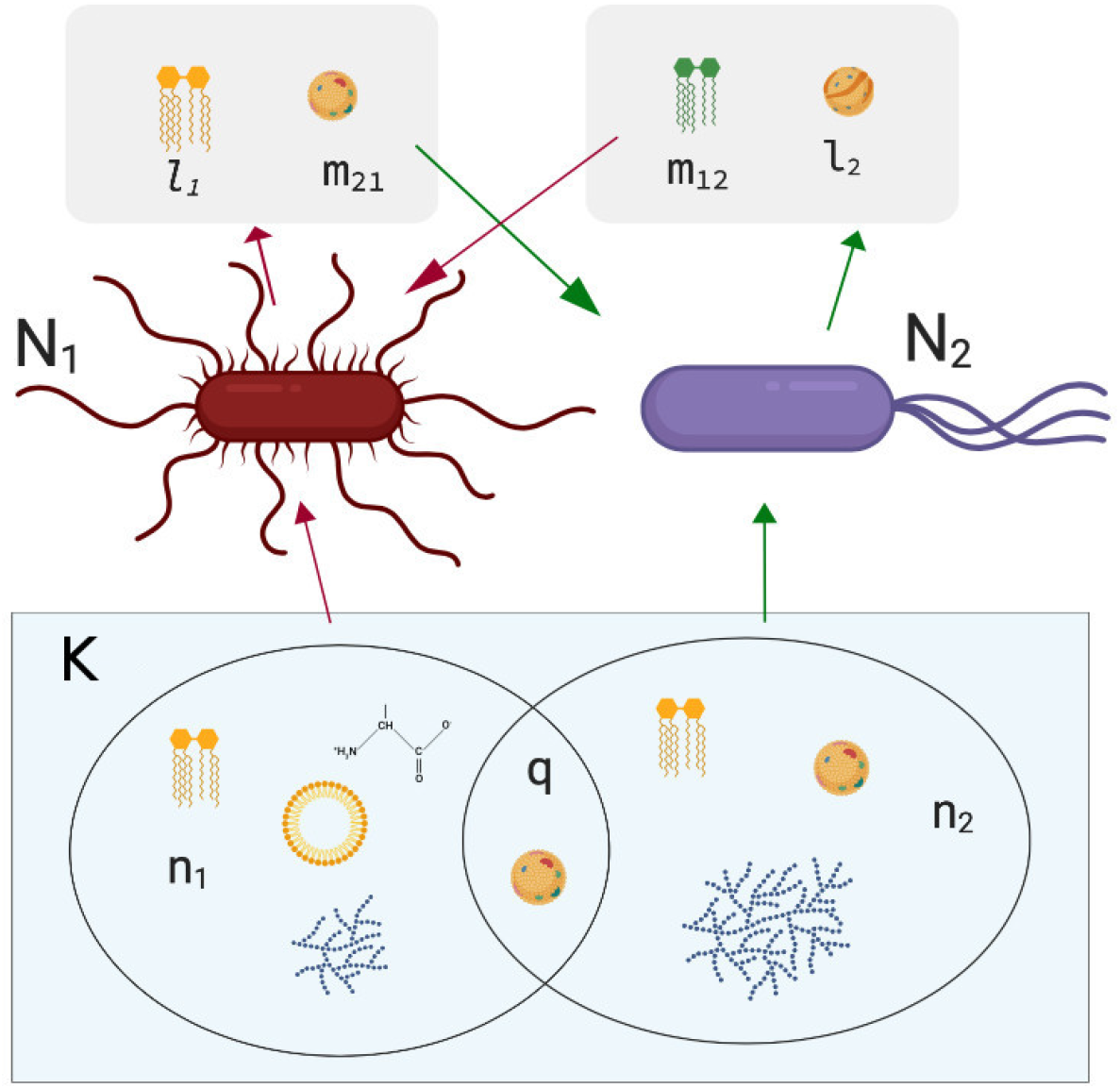
Sketch of the model for two microbial species. Figure created with ***BioRender (2019***).

### Adaptive dynamics

According to ***Equation 1***, the dynamics of species *i*’s population fits the pattern

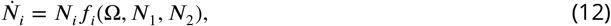

where Ω denotes the set of parameters {*u*_1_, *u*_2_, *w, α*_1_, *α*_2_}. The function *f*_*i*_(Ω, *N*_1_, *N*_2_) describes the per-capita growth rate, or *fitness*, of species *i*, a magnitude that depends on the population growth parameters as well as the population sizes of the two species involved. Any steady state of the community, 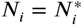, is defined by the equations

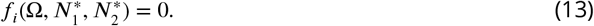

When a community is in a given steady state, a mutant may appear in one of the populations. If the mutant belongs to species *i*, its fitness will be a function 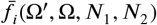, where the prime parameters are those of the mutant. If this fitness is negative the mutant will go extinct, otherwise it can increase its frequency in the population and replace the original genotype.

Adaptive dynamics (AD) is a method to exploit this idea to devise a set of differential equations for the demographic parameters. Under the assumption that the parameters of the mutant are a small perturbation of those of the original genotype, ***Dieckmann and Law (1996***) derived these differential equations from the master equation of the underlying stochastic mutation-selection process.

The derivation of these so-called canonical equations of AD implicitly assumes that the demographic parameters can vary independently of each other, and that mutations have the same probability to increase or decrease these parameters by the same small amount. Neither of these two conditions are met in our model under the scheme of mutations of this system (increasing or decreasing the number of resources). One kind of mutations amounts to changing the status of a randomly chosen external resource, i.e., adding the resource if it is new, or dropping it if the species was already using it. This means that *n*_*i*_ → *n*_*i*_ +1 with a probability proportional to the number of new resources, and *n*_*i*_ → *n*_*i*_ − 1 with a probability proportional to the number of resources already in use. But at the same time, *q*, the number of common resources to both species, can increase, decrease, or remain unchanged. ***Table 2*** summarises all mutational scenarios and their corresponding probabilities.

**Table 2.** Set of mutations in the system that change the number of external resources and their corresponding probabilities. Here *n*_*i*_ are the number of external resources consumed by species *i, q* the amount of them common to both species, and *u*_*i*_ ≡ *n*_*i*_/*K, w* ≡ *q*/*K*.

| initial state: $n_1, q$ | | | initial state: $n_2, q$ | | |
| --- | --- | --- | --- | --- | --- |
| mutated state | probability |  | mutated state | probability |  |
| $n_1 + 1, \quad q + 1$ | $u_2 - w$ | | $n_2 + 1, \quad q + 1$ | $u_1 - w$ | |
| $n_1 + 1, \quad q$ | $1 - u_1 - u_2 + w$ | | $n_2 + 1, \quad q$ | $1 - u_1 - u_2 + w$ | |
| $n_1 - 1, \quad q$ | $u_1 - w$ | | $n_2 - 1, \quad q$ | $u_2 - w$ | |
| $n_1 - 1, \quad q - 1$ | $w$ | | $n_2 - 1, \quad q - 1$ | $w$ | |

#### Box 1.

**Generalised logistic model of population dynamics**

The idea behind the population model of ***Stucchi et al. (2020***) is to extend Velhurst’s logistic equations of populations

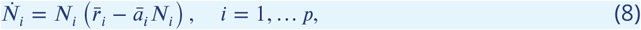

by making the parameters 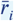 and *ā*_*i*_ to depend on the interactions with the environment as well as the population sizes of all species in the community as

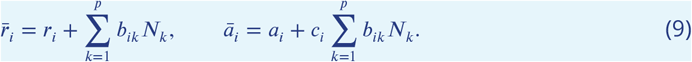

Here *r*_*i*_ is the vegetative growth rate of species *i, b*_*ik*_ is the rate of benefit (if positive) or hindrance (if negative) on species *i* due to the interaction with species *k*, and *p* is the total number of species in the ecosystem. The coefficients *a*_*i*_ measure intraspecific competitions (hence *a*_*i*_ > 0) due to a limitation of the environmental resources. As a matter of fact, in the standard Velhurst’s model 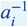 directly measures the carrying capacity of the environment. As of *c*_*i*_, the effect of these coefficients is better seen if we rewrite ***Equation 8*** and ***Equation 9*** as

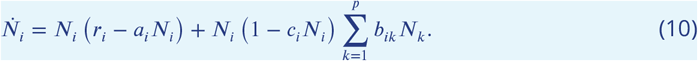

We can clearly see in this expression that choosing *c*_*i*_ > 0 induces a saturation on the interaction of the community with the focal species.

Incidentally, this generalised logistic model includes (through the coefficient *b*_*ii*_) extra terms of intraspecific interaction beyond the environmentally-induced, indirect competition accounted for by the −*a*_*i*_*N*_*i*_ term. Hence cooperation (*b*_*ii*_ > 0) or direct competition (*b*_*ii*_ < 0) can be readily incorporated to this model.

According to ***Stucchi et al. (2020***), the linear stability of the finite stationary solution for two species 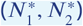 can be analyzed from the Jacobian matrix

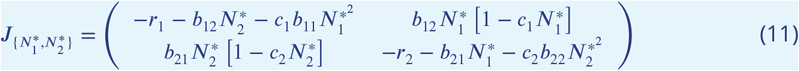

The second kind of mutations will change *m*_*ij*_, the amount of byproducts that species *i* take from species *j*, by ±1. As *α*_*i*_ is a multiplicative factor,

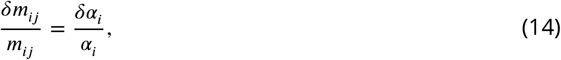

so *δm*_*ij*_ = ±1 implies

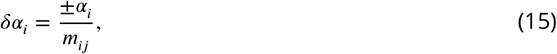

or, using ***Equation 4***,

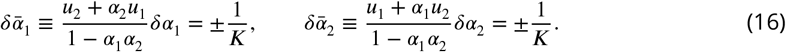

On the other hand, *δα*_*i*_ < 0 with probability *m*_*ij*_ /*m*_*i*_ = *α*_*i*_/*γ* and *δα*_*i*_ > 0 with probability *ℓ*_*i*_/*m*_*i*_ = (*γ* − *α*_*i*_)/*γ*.

Reconstructing the procedure of ***Dieckmann and Law (1996***) to obtain the canonical equation of AD, we arrive at a system of differential equations for *u*_1_, *u*_2_, *w, α*_1_, *α*_2_ (c.f. ***Equation 22*** and ***Equation 26*** in ***Appendix 1***; see the Supplementary Material for a more detailed derivation of the equations). Notice that the stationary populations 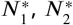 appear explicitly in the equations, and that these populations depend in turn on the parameters *u*_1_, *u*_2_, *w, α*_1_, *α*_2_.

### Evolutionary attractors

Although the differential system of ***Equation 22*** is very difficult to discuss analytically, in the many simulations that we have run using different sets of parameters and initial conditions the evolutionary attractors that we have observed all match just a few patterns. The triad of parameters (*u*_1_, *u*_2_, *w*) (fraction of resources consumed by each species and fraction of shared resources) is always found to end up in one of the three forms (1, *u, u*), (1 − *u, u*, 0), or (1 − *u, u*, min(1 − *u, u*)), with 0 ≤ *u* ≤ 1. If (1, *u, u*) is reached, the evolutionary attractor turns out to be either competition or antagonism. In particular, in the case (1, 1, 1) the attractor is always competition. As expected, when the final state is mutualism or commensalism no resources are shared (*w* = 0) (there is no need for competing for resources), so when (1 − *u, u*, 0) is reached the system ends up being mutualistic.

With respect to the cross-feeding efficiencies, *α*_*i*_, the system generally evolves towards extreme values of these parameters (*α*_*i*_ = 0 or *α*_*i*_ = *γ*). Only when the system evolves to competition can some intermediate values be found. As expected, cross-feeding efficiencies reach their maximum (*α*_1_ = *α*_2_ = *γ*) for mutualism and their minimum (*α*_1_ = *α*_2_ = 0) mainly for competition.

It is worth mentioning that all kinds of mutualisms are found to be evolutionary attractors for some initial conditions. Another relevant observation is that evolution sometimes drives ecosystems to extinction. This is no longer a surprise because it is a result that has already been empirically observed (e.g. in viruses (***Turner and Chao, 1999***)), but the idea that evolution can degrade ecosystems would have shocked evolutionists of the 19th and early 20th century because it contradicts the notion of evolution as a sort of ‘optimiser’.

### Evolutionary transitions between types of ecological interactions

In order to illustrate the kind of evolutionary transitions between types of ecological interactions that this system can produce, we performed an exhaustive exploration of parameters. We fixed the environmental parameters *a*_1_ = *a*_2_ = *c*_1_ = *c*_2_ = 0.001 and, without loss of generality, chose *r*_10_ = 1 (this simply sets the evolutionary time scale). For the other species, we explored uniformly the interval 0 < *r*_20_ ≤ 1 and then zoomed in the region 0 < *r*_20_ ≪ *r*_10_ by sampling the interval 0 < *r*_20_ ≤ 0.1. For *b*_*i*0_ we sampled uniformly the range 0 ≤ *b*_*i*0_ ≤ 0.01 and then zoomed in the interval 0 ≤ *b*_*i*0_ ≤ 0.001. Likewise, nine different, uniformly spaced values of the metabolic efficiency *γ* within the range 0 < *γ* < 1 were explored.

For each set of parameters we generated random initial conditions for *u*_*i*_, *w*, and *α*_*i*_ within the ranges 0 ≤ *u*_*i*_ ≤ 0.99, 0 ≤ *w* ≤ *min*(*u*_1_, *u*_2_), and 0 ≤ *α*_*i*_ ≤ *γ*, and kept only those that generated viable populations. Then we let each of these remaining cases evolve according to ***Equation 22*** and ***Equation 26***. We recorded 1000 runs that resulted in viable populations for each set of parameters, discarding all initial conditions that led to no viable stationary populations but keeping track of those that eventually led to extinction. Notice that the number of resources or the mutation probability are only relevant to set the evolutionary time scale (see ***Appendix 1***).

In order to catalogue the resulting evolutionary attractors we have followed the classification of ***Table 1***—considering a state as commensalist if one of the parameters *b*_*ij*_ is positive and the other one is smaller than 10^−8^.

The results of this numerical study of the model are summarized in ***Figure 2, Figure 3, Figure 4***, and ***Figure 5*** (for the distribution of the initial and final states see the Supplementary Material). In what follows we describe in more detail the transitions from an initial ecological state to another one that we observed, depending on the choice of parameters *r*_*i*0_ and *b*_*i*0_.

**Figure 2.**
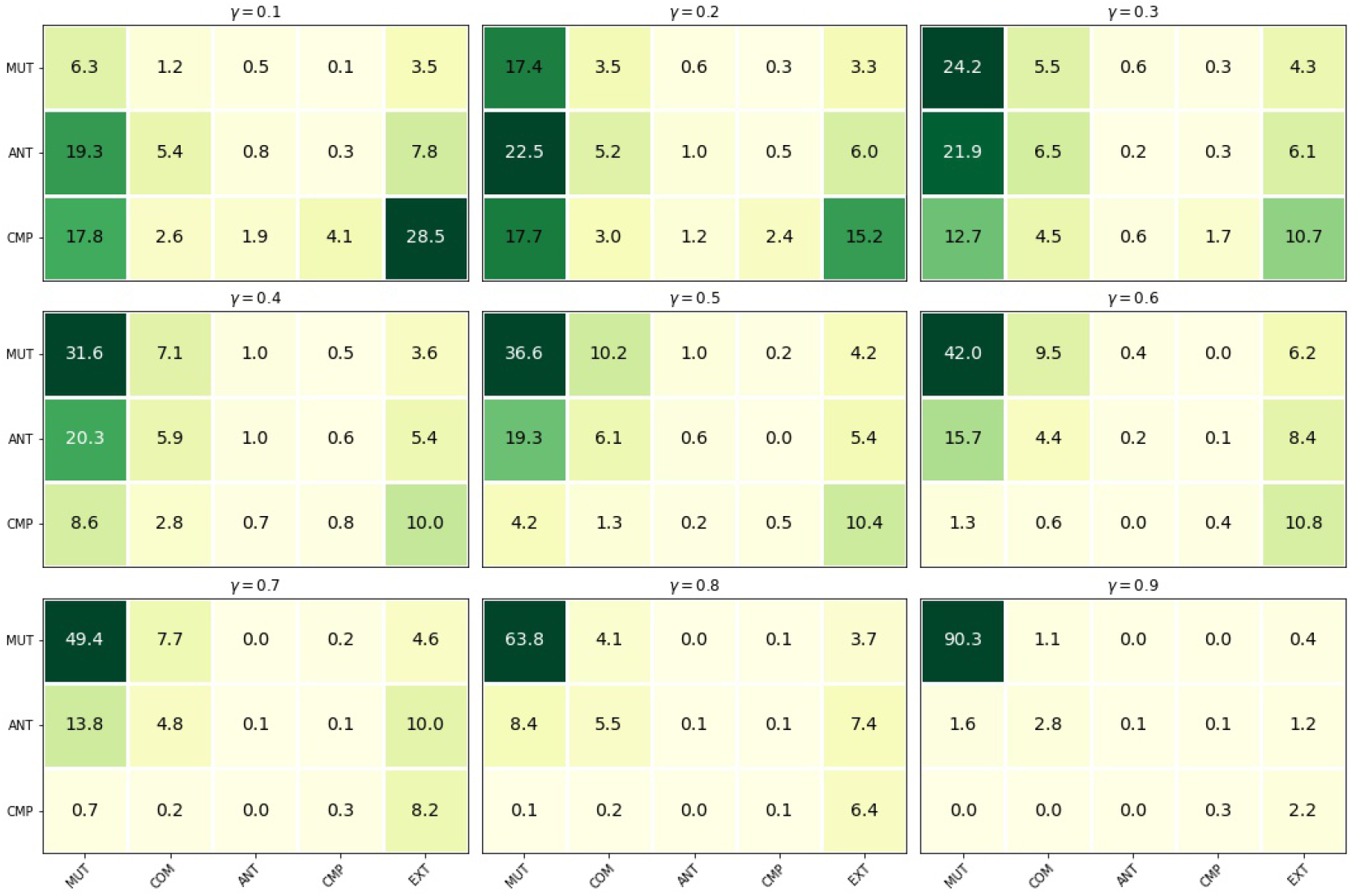
Evolutionary transitions for values of *γ* ranging from 0.1 to 0.9. In rows, we show the initial ecological interaction states (mutualism (MUT), commensalism (COM), antagonism (ANT), and competition (CMP)), and in columns, the final ecological interaction states—with the addition of extinction (EXT). Figures denote the probability of the corresponding transitions. Parameter values: *r*_20_ ≤ 1 and *b*_*i*0_ ≤ 0.01. For *γ* ≤ 0.2, the initial states are mainly ANT and CMP, and they mostly end up in MUT or EXT. For *γ* ≥ 0.3, the initial states are mainly MUT and remain MUT.

**Figure 3.**
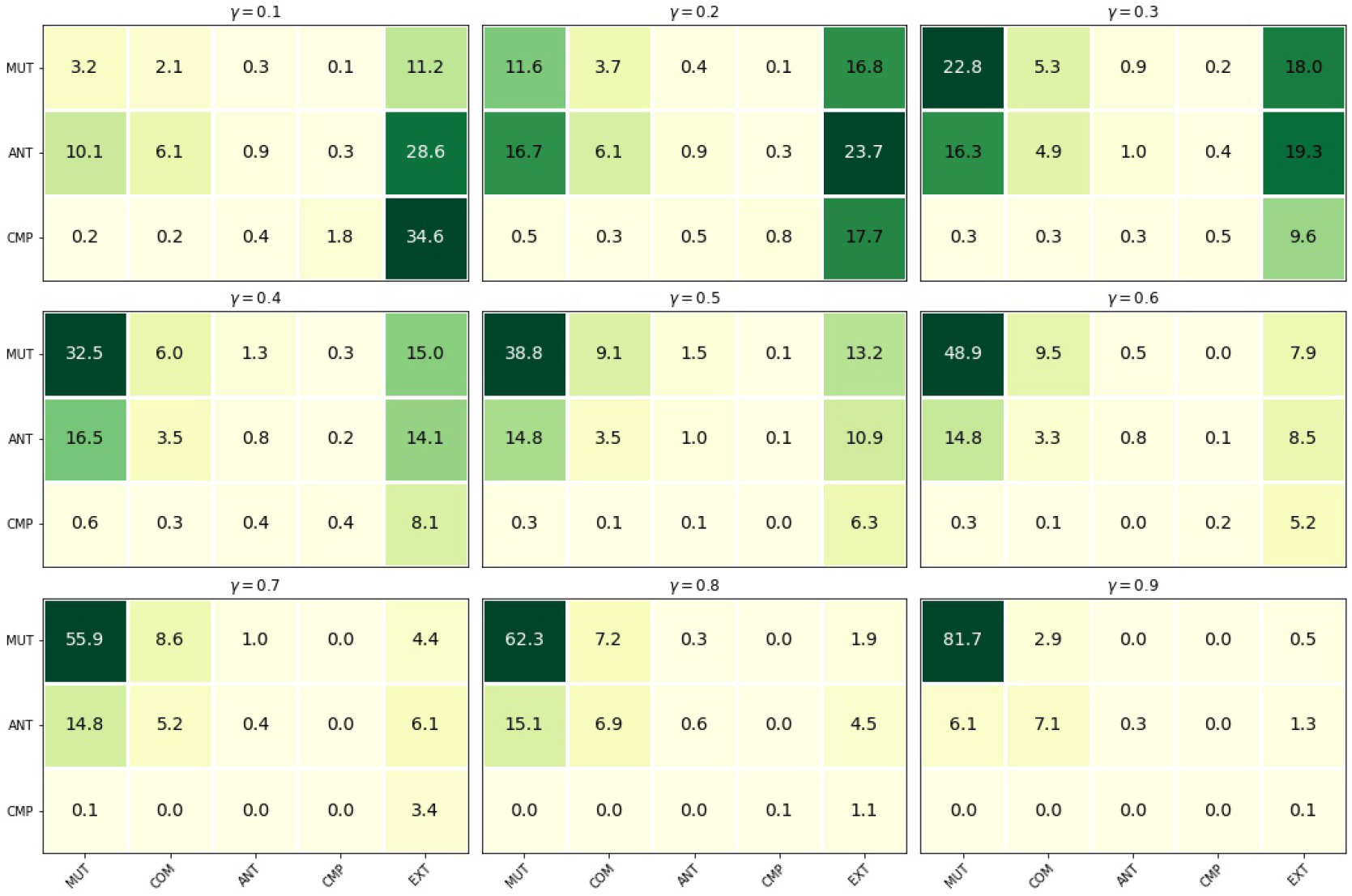
As ***Figure 2***, for parameters values *r*_20_ ≤ 0.1 and *b*_*i*0_ ≤ 0.01. For *γ* ≤ 0.2, the initial states are mainly CMP and ANT, although they mostly end up in EXT. For *γ* ≥ 0.3, the initial states are mainly MUT and remain MUT.

**Figure 4.**
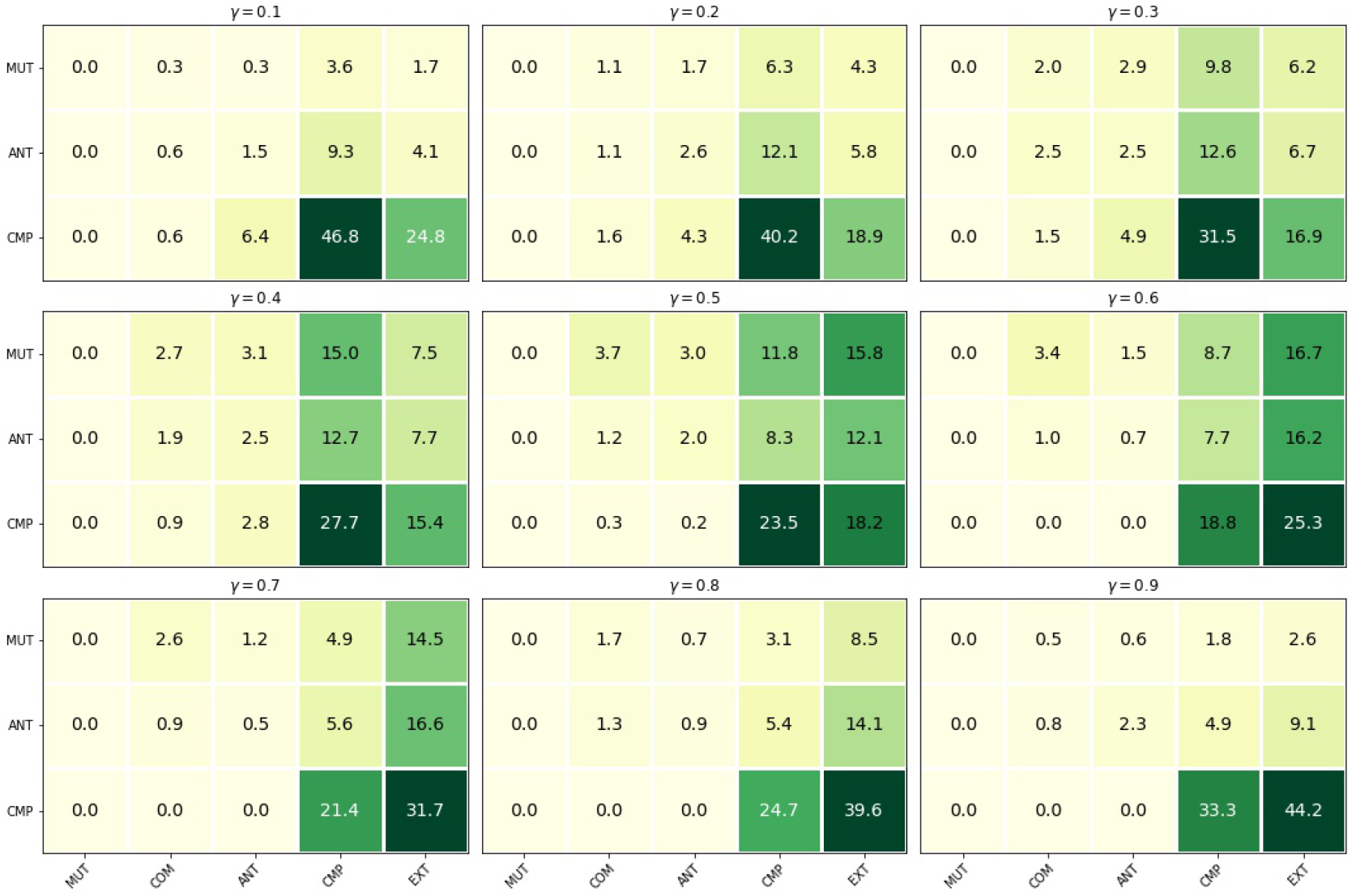
As ***Figure 2***, for parameters values *r*_20_ ≤ 1.0 and *b*_*i*0_ ≤ 0.001. For *γ* ≤ 0.3, the initial states are mainly CMP, and they mostly stay as CMP. For 0.3 ≤ *γ* ≤ 0.7, communities can also begin as MUT or ANT. For *γ* ≥ 0.6, most populations end up in CMP or EXT.

**Figure 5.**
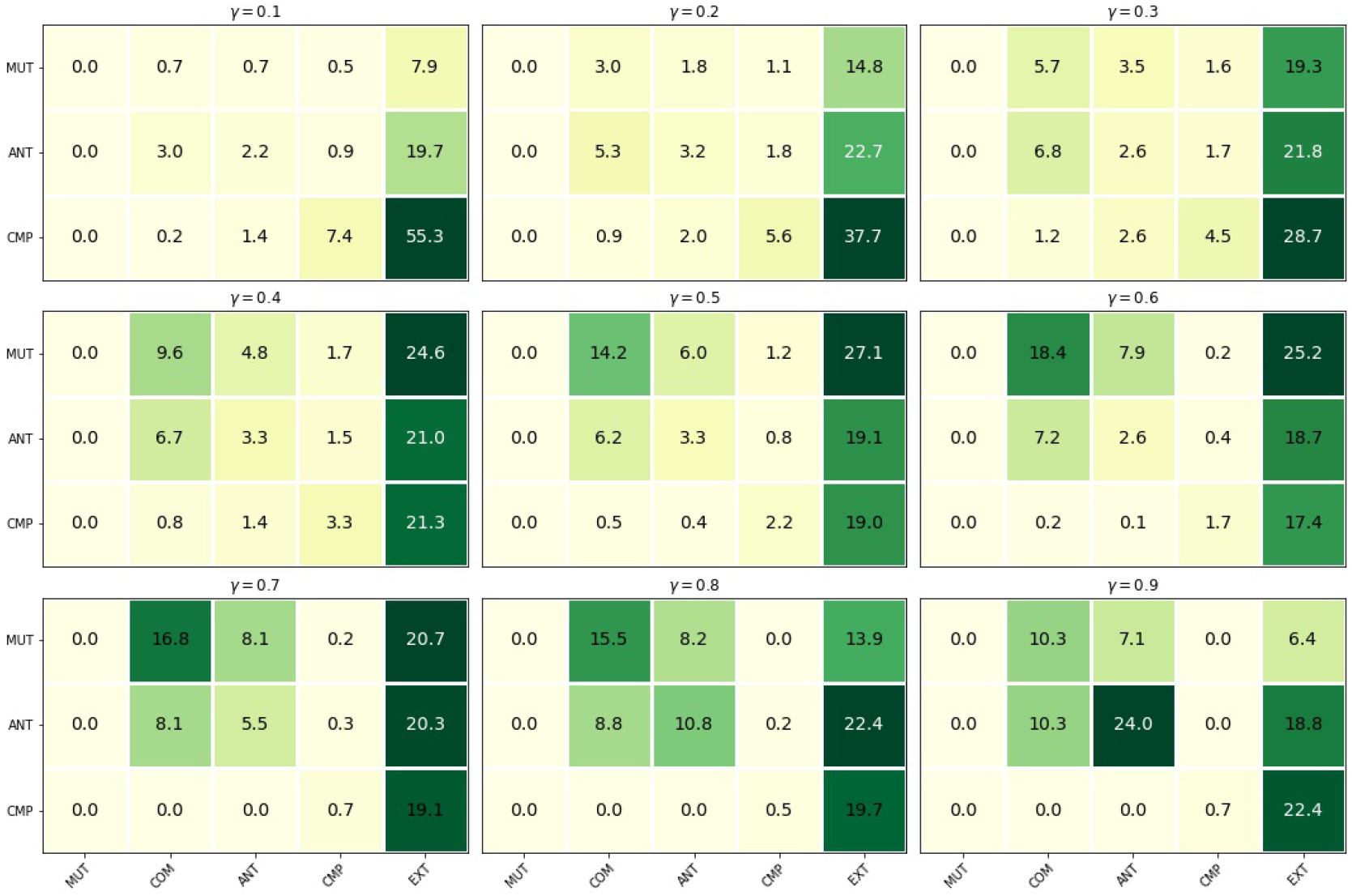
As ***Figure 2***, for parameters values *r*_20_ ≤ 0.1 and *b*_*i*0_ ≤ 0.001. For *γ* ≤ 0.3, the initial states are predominantly CMP, and they mostly end in EXT. For *γ* ≥ 0.4, the initial states are predominantly MUT, although EXT continues to be the most common evolutionary fate. However, for *γ* ≥ 0.8, COM and ANT become relatively common ending states.

*r*_20_ ≤ 1, *b*_*i*0_ ≤ 0.01

(See ***Figure 2***.) For this choice of parameters most evolutionary pathways ended up in mutualism. Depending on the value of *γ*, the distribution of initial and final states was like this:

***γ*** = **0.1:** 90% of the initial states were antagonistic or competitive, whereas the final states mostly distributed between mutualism (∼ 40%) and extinction (∼ 40%).

***γ*** = **0.3:** The initial states were competition, antagonism, and mutualism with approximately the same probability, and the final states were mostly mutualism (∼ 60%) and extinction (∼ 20%). Facultative mutualism was dominant in this case.

***γ*** = **0.5:** The initial states included collaborative or detrimental relationships almost equally (in nearly all cases mutualism was obligate for one species). The final states were mostly mutualistic (60%), and only 2% were antagonistic and 1% competitive (the remaining evolutionary pathways ended up in commensalism or extinction).

***γ*** = **0.7:** Most initial states were obligate mutualism for both species (21%), obligate mutualism for only one species (33%), and facultative mutualism (8%). Final states were essentially obligate mutualism for both species (62%) or extinction (23%).

***γ*** = **0.9:** 89% of the initial states were obligate mutualism for both species that ended up in the same final states (no transition).

*r*_20_ ≤ 0.1, *b*_*i*0_ ≤ 0.01

(See ***Figure 3***.) Most evolutionary pathways ended up in mutualism or went extinct. Depending on the value of *γ*, the distribution of initial and final states was like this:

***γ*** = **0.1:** 63% of the initial states were antagonistic or competitive, but only 4% of the cases remained as such. Of the rest, 14% became mutualistic and 74% went to extinction.

***γ*** = **0.2:** The initial states were mainly antagonistic (66%) or mutualistic (33%), whereas those final states which did not go to extinction (58%) were generally mutualistic (29%).

***γ*** = **0.6:** Initially mutualistic states (∼ 60%) mostly remained mutualistic, whereas intially antagonistic states (38%) evolved to either commensalism or extinction.

***γ*** > **0.6:** Initial and final states were mainly mutualistic.

*r*_20_ ≤ 1, *b*_*i*0_ ≤ 0.001

(See ***Figure 4***.) For this parameters the system evolved mainly to competition or went extinct. With such a small interaction parameters *b*_*i*0_ mutualism is dramatically hindered—even in the cases where one third of the initial states were mutualistic (for *γ* = 0.5). More than 40% of the system started in competition and remained as such or went extinct—the proportion of which changed upon increasing *γ* from 46.8% competition vs. 24.8% extinction, to 33.3% competition vs. 44.2% extinction.

*r*_20_ ≤ 0.1, *b*_*i*0_ ≤ 0.001

(See ***Figure 5***.) As in the previous case, the system cannot evolve into a mutualistic state. For different values of *γ* we found:

***γ*** = **0.1:** 63% of the initial states were competition, and only 9% of them remained as competition— the rest went extinct. Overall, 83% of the initial states ended up extinct.

***γ*** = **0.5:** Two thirds of the initial states went extinct. The rest of them ended up mostly in commensalism (20–25%).

***γ*** > **0.5:** Overall, more than 47% evolutionary pathways ended up in extinction. For *γ* = 0.9, 24% of them started and finished in antagonism.

### Transient states

Because of the rich evolutionary dynamics of this model, evolutionary transitions are not the only relevant feature to focus on. Particular sequences of transient ecological states along evolutionary pathways are as interesting—if not more so. In ***Figure 6, Figure 7, Figure 8, Figure 9, Figure 10***, and ***Figure 11*** we show the time evolution of the demographic parameters of just a few examples, chosen because of their peculiar sequence of intermediate transitions or because they resemble actual transitions observed in nature. (For the time evolution of the stationary populations *N*_*i*_ and their evolutionary parameters, see the Supplementary Material).

**Figure 6.**
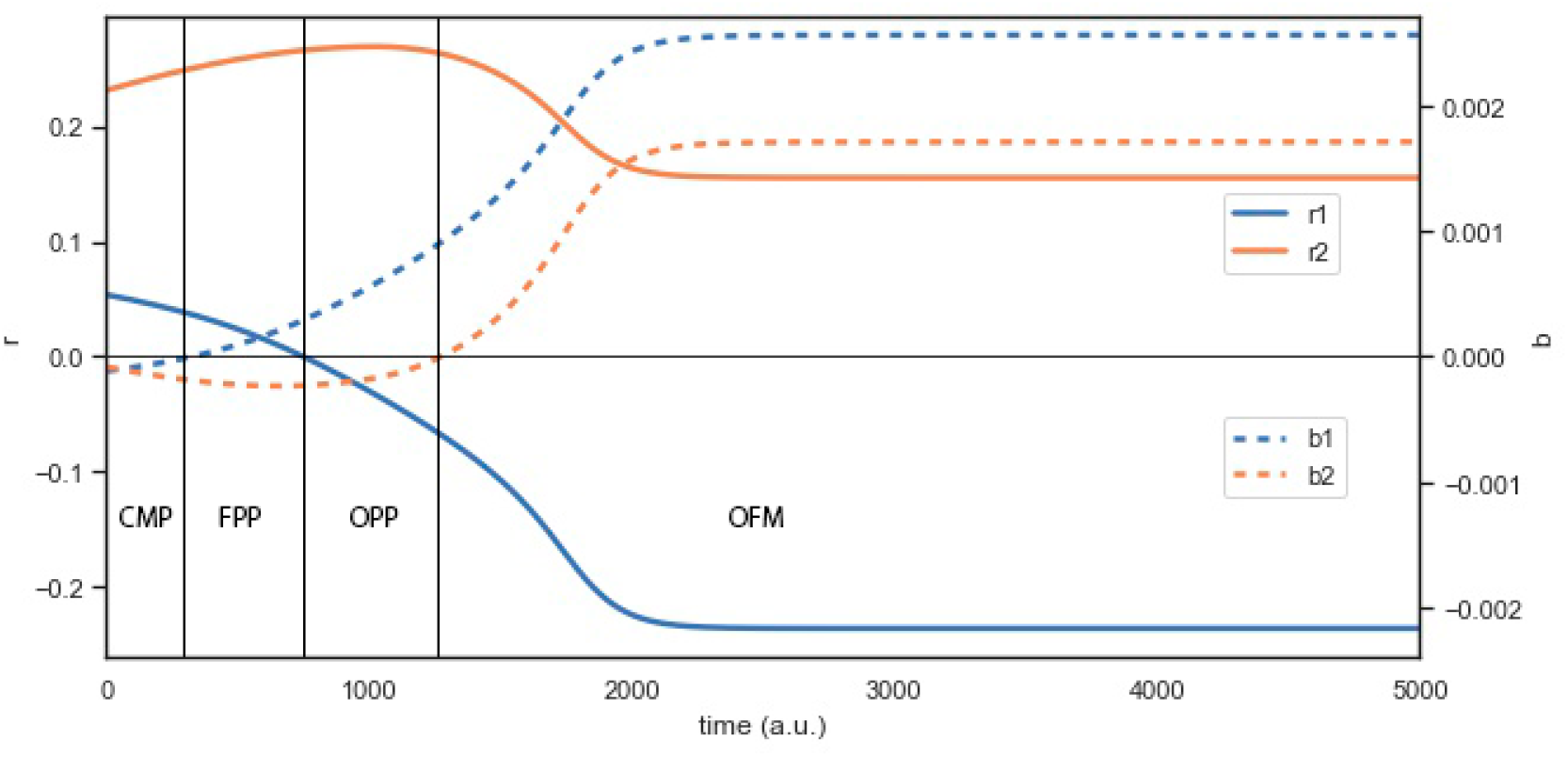
Evolutionary transition of the population parameters *r*_*i*_ and *b*_*i*_ of two species. The community starts in competition (CMP) and ends in obligate-facultative mutualism (OFM), going through intermediate states of facultative parasitism (FPP) and obligate parasitism (OPP). Initial parameters: *r*_01_ = 1, *r*_02_ = 0.65727, *b*_01_ = 0.00414, *b*_02_ = 0.00447, *u*_1_ = 0.17967, *u*_2_ = 0.78500, *w* = 0.10050, *α*_1_ = 0.08365, *α*_2_ = 0.32459, *γ* = 0.5.

**Figure 7.**
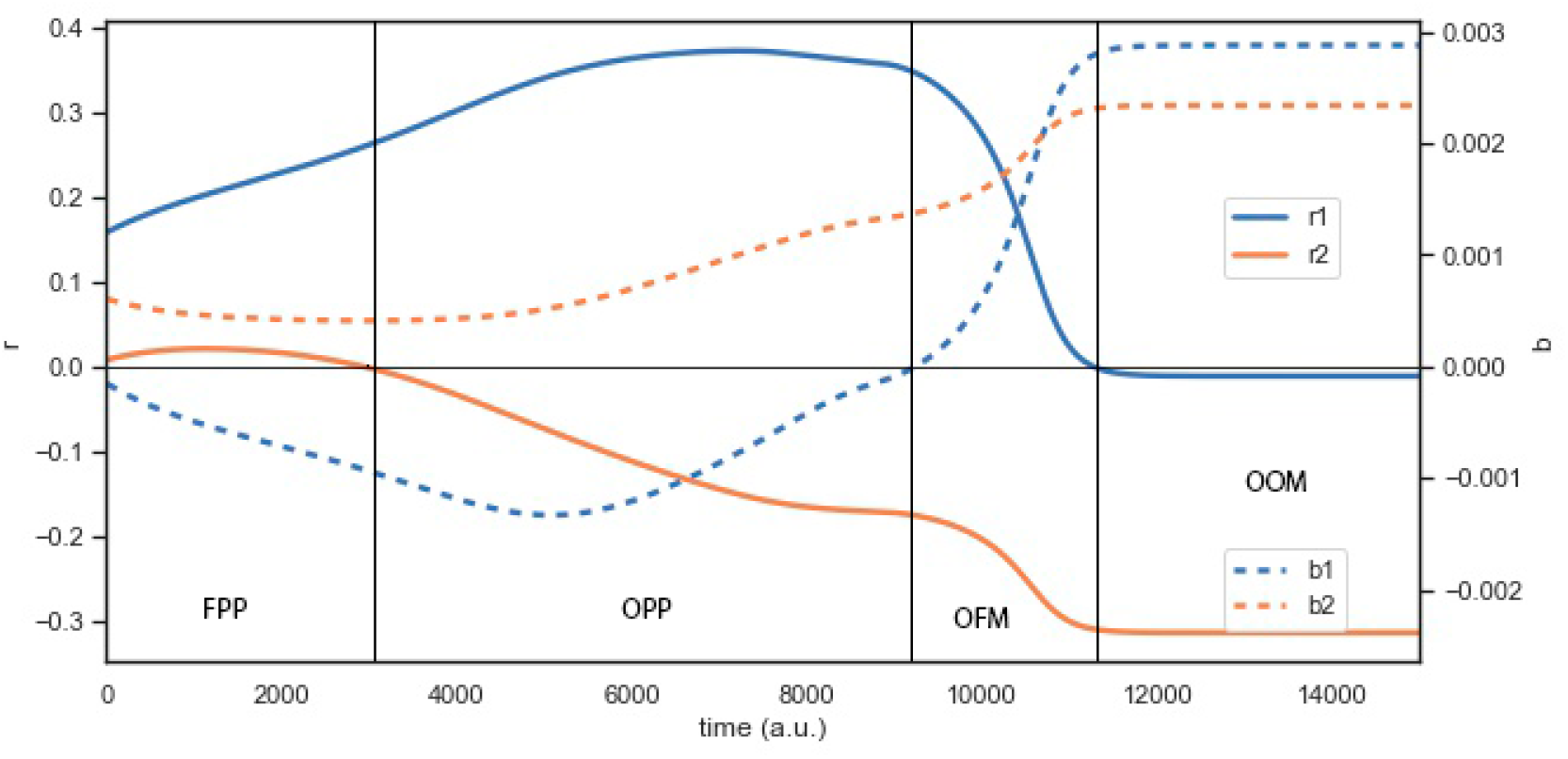
Evolutionary transition of the population parameters *r*_*i*_ and *b*_*i*_ of two species from facultative parasitism (FPP) to obligate-obligate mutualism (OOM), with intermediate states of obligate parasitism (OPP) and obligate-facultative mutualism (OFM). Initial parameters: *r*_01_ = 1.0, *r*_02_ = 0.64396, *b*_01_ = 0.00474, *b*_02_ = 0.00262, *u*_1_ = 0.60097, *u*_2_ = 0.62973, *w* = 0.16897, *α*_1_ = 0.13282, *α*_2_ = 0.54215 and *γ* = 0.6.

**Figure 8.**
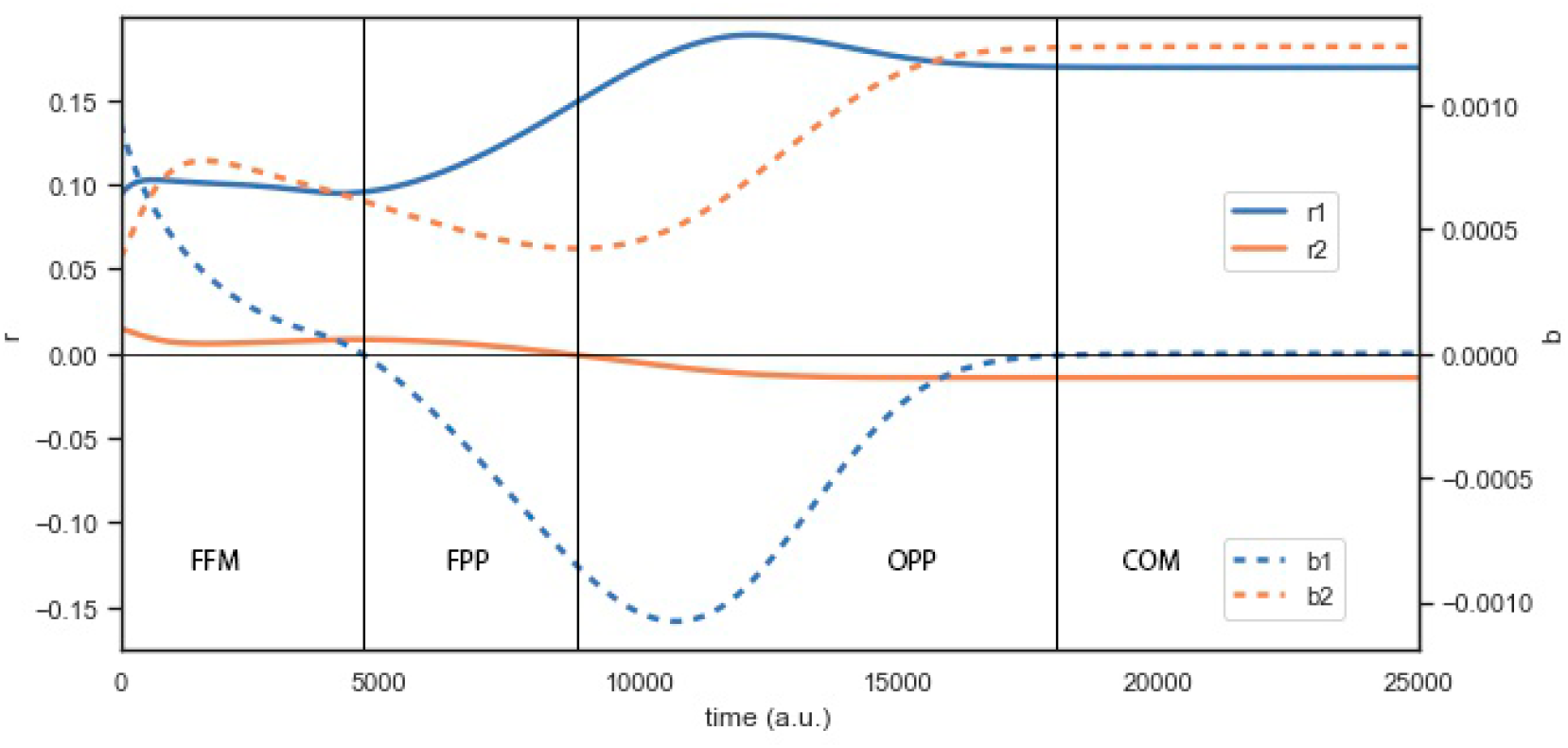
Evolutionary transition of the population parameters *r*_*i*_ and *b*_*i*_ of two species from facultative-facultative mutualism (FFM) to commensalism (COM), with intermediate states of facultative parasitism (FPP) and obligate parasitism (OPP). Initial parameters: *r*_01_ = 1.0, *r*_02_ = 0.09827, *b*_01_ = 0.00463, *b*_02_ = 0.00313, *u*_1_ = 0.79063, *u*_2_ = 0.80318, *w* = 0.00460, *α*_1_ = 0.22106, *α*_2_ = 0.12717, and *γ* = 0.7.

**Figure 9.**
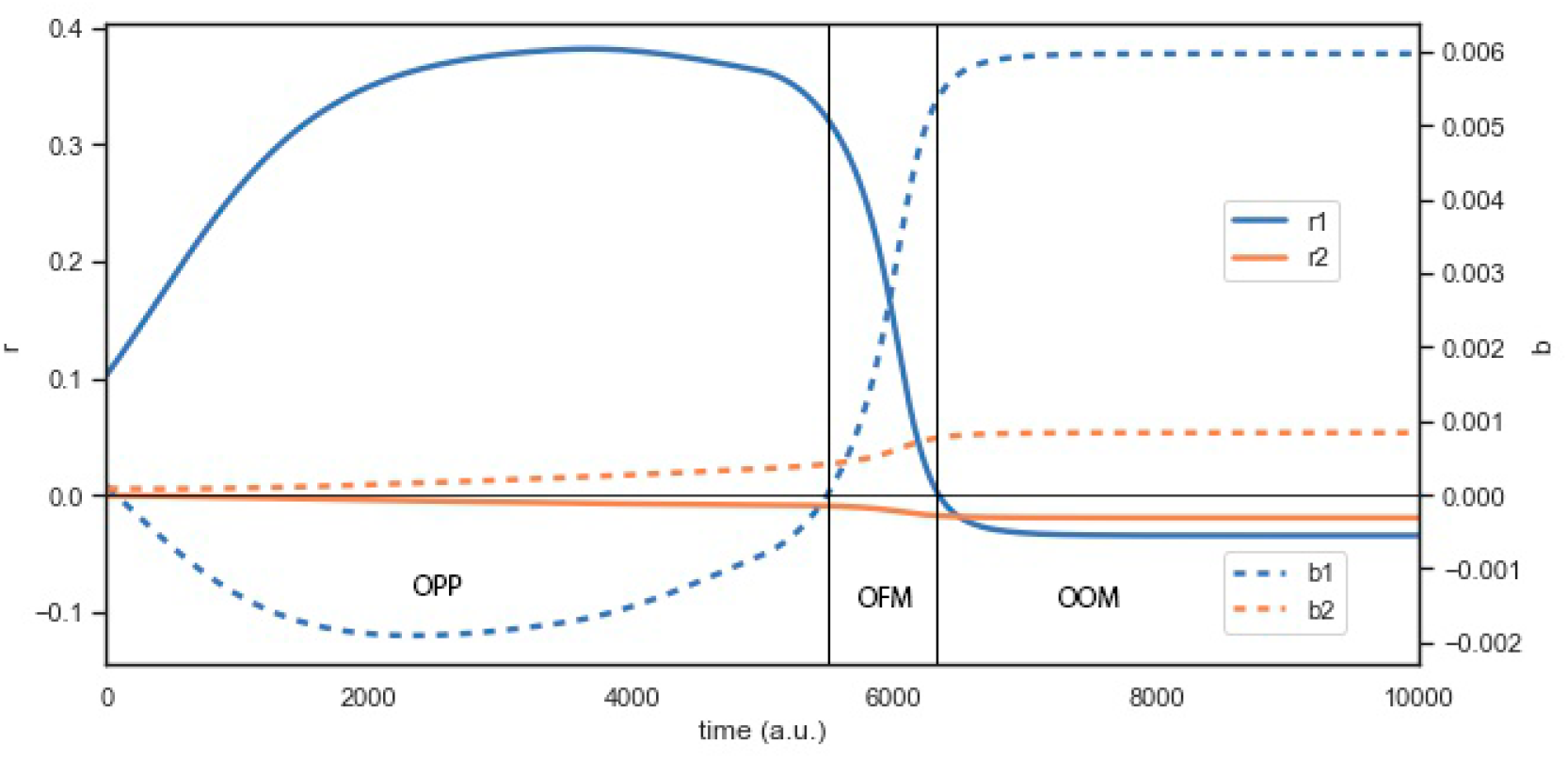
Evolutionary transition of the population parameters *r*_*i*_ and *b*_*i*_ of two species from obligate parasitism (OPP) to obligate-obligate mutualism (OOM), with an intermediate state of obligate-facultative mutualism (OFM). The initial OPP has a long stasis, wherear the intermediate OFM is much shorter in evolutionary scale. Initial parameters: *r*_01_ = 1, *r*_02_ = 0.04230, *b*_01_ = 0.00962, *b*_02_ = 0.00096, *u*_1_ = 0.45854, *u*_2_ = 0.26628, *w* = 0.12217, *α*_1_ = 0.28321, *α*_2_ = 0.35339, and *γ* = 0.6.

**Figure 10.**
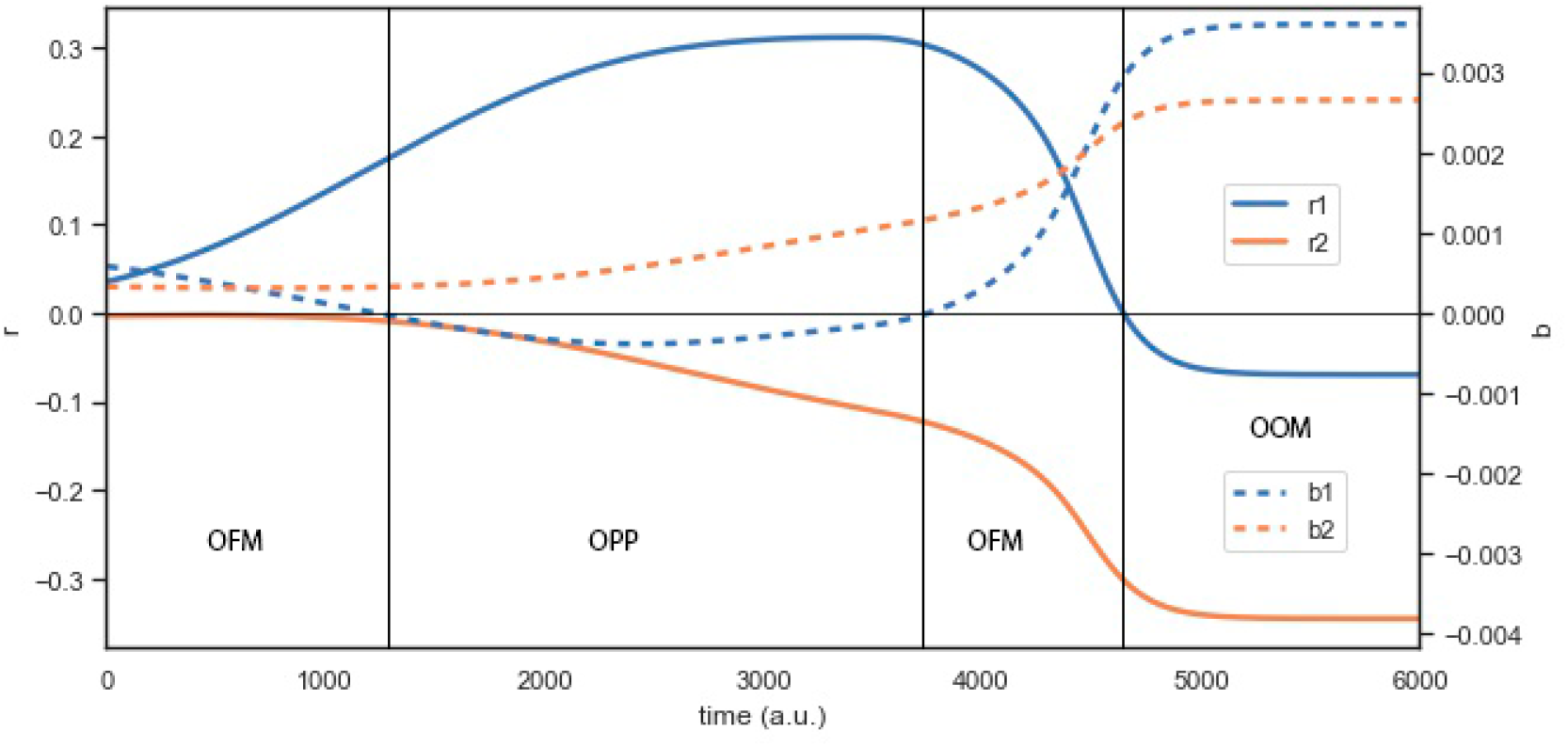
Evolutionary transition of the population parameters *r*_*i*_ and *b*_*i*_ of two species from obligate-facultative mutualism (OFM) to obligate-obligate mutualism (OOM), with an intermediate state of obligate parasitism (OPP) and obligate-facultative mutualism (OFM). Initial parameters: *r*_01_ = 1.0, *r*_02_ = 0.80306, *b*_01_ = 0.00561, *b*_02_ = 0.00310, *u*_1_ = 0.26642, *u*_2_ = 0.17242, *w* = 0.01485, *α*_1_ = 0.40480, *α*_2_ = 0.31492, and *γ* = 0.6.

**Figure 11.**
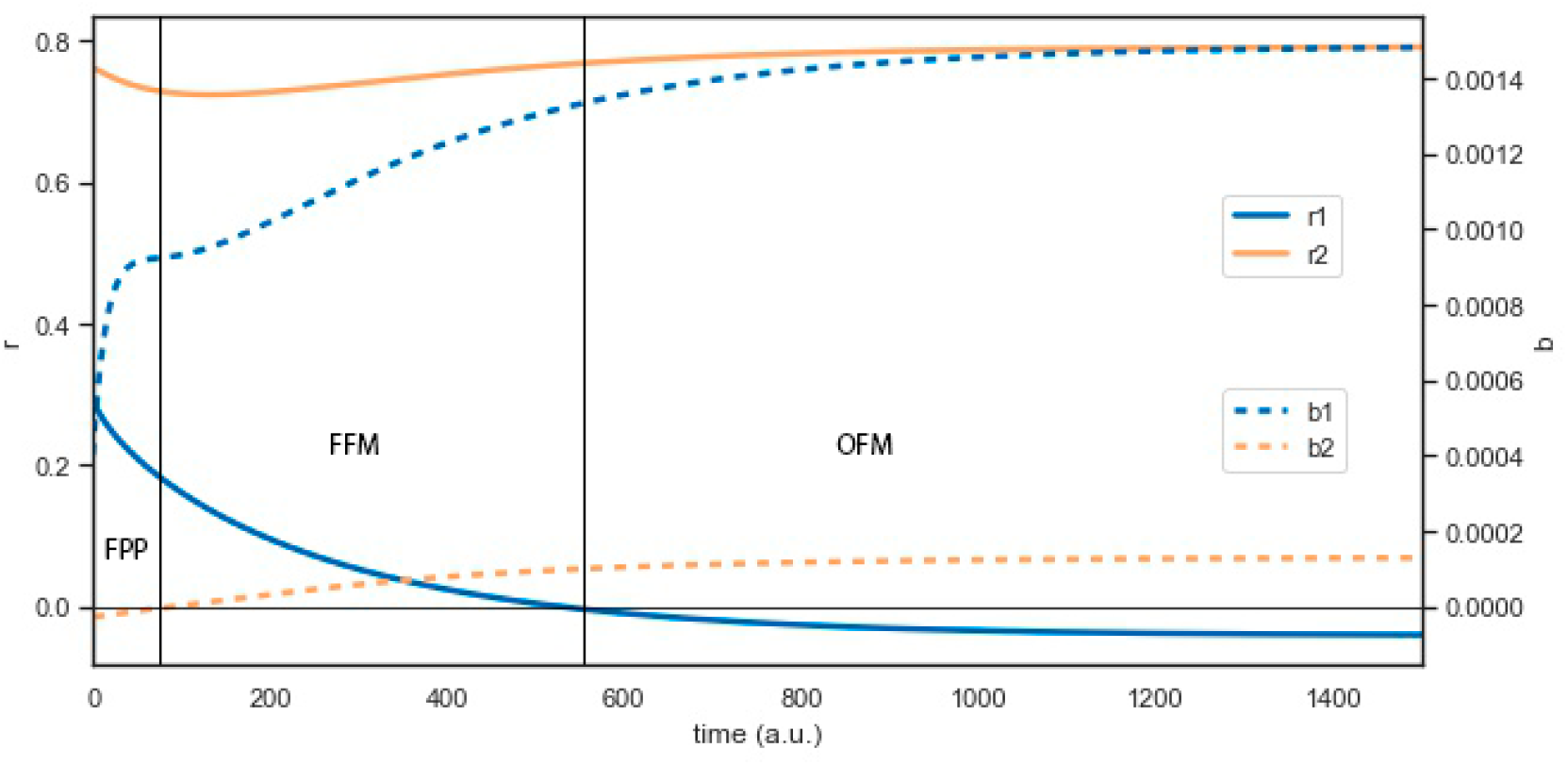
Evolutionary transition of the population parameters *r*_*i*_ and *b*_*i*_ of two species from facultative parasitism (FPP) to obligate-facultative mutualism (OFM), with an intermediate state of facultative-facultative mutualism (FFM). Initial parameters: *r*_01_ = 1.0, *r*_02_ = 1.0, *b*_01_ = 0.00713, *b*_02_ = 0.00309, *u*_1_ = 0.38958, *u*_2_ = 0.96352, *w* = 0.05325, *α*_1_ = 0.10848, *α*_2_ = 0.08866, and *γ* = 0.2.

***Figure 6*** shows two species with positive vegetative growth rate that are initially in competition. Over time one of them starts parasitizing the other until becoming dependent on it. Eventually, the other species ‘learns’ to take advantage of the parasite and the relation ends up as a mutualism.

In ***Figure 7*** species 2 starts as a parasite of species 1, although parasitism is facultative. Soon the parasite becomes dependent, and this situation remains like this for a while, until suddenly (in evolutionary terms) the relation evolves into a mutualism, facultative at first for species 1, but eventually obligate for both species.

A similar co-evolutionary pathway can be found in nature between Macrotermitinae (species 1) and fungi (species 2). According to ***Nobre et al. (2010***) and ***Aanen et al. (2002***), both the fungi and the fungus-growing termites are obligate mutualists since they need each other in order to survive and reproduce. As stated by ***Margulis and Sagan (2002***), it is plausible that the origin of the termite-fungi mutualistic relation was an infection (a specialized infestation) of the termites guts with fungi spores, which led them to defend themselves by domesticating the fungi, controlling and limiting their growth.

***Figure 8*** is an example of an opposite transition, in which one of the species of an initially mutualistic system evolves into a parasite of the other (first facultative, eventually obligate) which, over time, develops a commensalistic relation with the parasitized species.

Commensalism arises frequently in nature. For instance some algae and Ascomycota fungi do not form lichens, but descend of lichen-forming fungi ancestors. Non-lichenized fungi might obtain their nutrients acting as commensalists of lichen-forming fungi and algae (***Lutzoni et al., 2001***). Although the evolution of the lichen symbiosis is believed to have appeared multiple independent times (***Divakar et al., 2015***), no specific route for this formation has been described with certainty, to our knowledge.

There are many other cases of known mutualist relations that had become commensalistic. Zooxanthellae and octocorals form an ancestral facultative mutualistic relation where the dinoflagellates contribute to the energy budget of the invertebrates, by being host inside them. However, ***Oppen et al. (2005***) showed that there is compelling evidence of a FFM→COM transition, within the octocoral family Melithaeidae. Zooxanthellae and antheopleura form a similar relation and ***Geller and Walton (2001***) stated that many species of the sea anemones seem to have lost the mutualistic relationship with the algae. *Gymnodinium* algae provide an energetic supply to many marine invertebrates, which in return act as protective hosts. ***Wilcox (1998***) show, through a molecular phylogenetic analysis, that *Gymnodinium* is a genus that incorporate both mutualistic and independent species even though they all descend from a common symbiotic ancestor.

In ***Figure 9*** an initially parasitic relation, with a long period of stasis, eventually evolves into an interdependent mutualistic relation after crossing a brief period of facultative mutualism. Notice that the co-evolution of *Legionella jeonii* and *Amoeba proteus* described in the Introduction illustrates this kind of evolutionary pathway (***Jeon and Lorch, 1967***).

Another example, ***Figure 10***, exhibits a case that begins and ends in a mutualistic state, but not before going through a period of parasitism. This case reveals that, even though some of the cases reported in the statistics of the previous section appear not to have undergone any transition whatsoever, they nevertheless can come across different intermediate states before reaching their evolutionary stable state.

As a last example, ***Figure 11*** exhibits a case that begins in facultative parasitism and ends in a obligate-facultative mutualism, going through a period of facultative-facultative mutualism. This case is similar to the case of ants and aphids from several genera, whose relations have gone through all mutualistic and commensalistic nuances, as we pointed out in the Introduction.

Attine ants and fungi might have also undergone a similar pathway, since they form mutualistic relations that range from almost mutually obligatory to facultative—at least for the fungi. In particular, ***Nobre et al. (2010***) indicate that the fungal symbionts of the higher attines depend almost exclusively on the ants—although occasionally they might reproduce sexually. However, ***Schultz and Brady (2008***) and ***Currie et al. (2003***) also show that fungi are capable of living without the symbiotic relation with the ants.

Several hypotheses have been drawn to explain the evolutionary origin of the attine ant-fungus relation. According to ***Mueller et al. (2001***), even though it is widely accepted that fungi were part of the ancestral ant diet (which would constitute a predatory relation), neutral coexistence, where ants acted as accidental vehicles of fungal dispersion, may be a more viable explanation (in which case, the ancestral relation would be commensalistic, instead of predatory).

## Discussion

Admittedly, the model we have proposed and analyzed in this paper is a far cry from any realistic description of microbial interactions. For instance, we have deliberately ignored any detail on the metabolic processes involved—which are determinant in deciding which products can or cannot be re-used—or kept the environmental conditions constant—hence neglecting any effect that a change in the environment might bring into the interactions (***Thompson, 1988; Hernández, 1998***). For these and many other drastic simplifications we have assumed in its design, it is only fair to call our model a ‘toy model’. And yet this is precisely one of its virtues, because what this model makes clear is that very few assumptions about the way two species can interact leads to drastic changes in the ecological scenario. More and more, empirical evidence is showing that evolutionary transitions in the ecological interactions between species of a community are the norm, rather than the exception. The model proposed here provides a proof of concept in this sense, because it links natural microscopic interactions between microbes to subsequent changes in their ecological relationships.

With a simple bookkeeping of the amount of resources that are taken from the environment, of those that are excreted as byproducts of metabolic reactions, and of the amount of byproducts from the other species that can be re-used for their own purpose, the model can cover virtually all possible ecological interactions between two species, and show, using adaptive dynamics, that these interactions evolve, going through different scenarios until reaching a final stable state. The model also provides some clues about general trends that can occur in real situations. For instance, there is a marked trend toward the emergence of mutualistic interactions, even in systems that start in competition or show antagonistic relationships. Also, many of the observed pathways have a real counterpart, because similar ones have been documented for actual species (microbial or otherwise).

The model relies upon the availability of a phenomenological model of ecological interactions (***García-Algarra et al., 2014; Stucchi et al., 2020***) that is capable of describing mutualistic, competitive, and antagonistic interactions with a simple tuning of the parameters—in a way that generalized Lotka-Volterra models are not capable of. The clue in devising our present evolutionary model has been linking those parameters to microscopic interactions between the species involved— which we have chosen as microbes for the sake of simplicity.

There are at least two ways in which this work can be extended. One is making more detailed and realistic assumptions on the microscopic interactions that occur between the two species and see whether the trends observed in this toy model are or are not confirmed. Obviously, introducing further details will impose constraints that the present model is currently free of. This constraints will bias the distribution of scenarios we observe in different ways, and will do so differently for different species—for which the details can vary from instance to instance. This will provide interesting information about the connection between microscopic mechanisms and ecological transitions.

The other way to extend this work is to consider more than two species. We can only imagine the richness of ecological scenarios that such an extension will reveal, even with a mechanism as simple as the straightforward bookkeeping of resources we have implemented. The computational complexities of this extension are evident by just looking at the involved calculations that only two species has led to. This is one of the reasons why we have decided to postpone such a study for future research—the other one being that just two species are enough to provide the proof of concept we aimed at with this work.

## Methods

Numerical simulations were performed in Python (***Stucchi and Pastor, 2021***). For the integration of the AD equations (***Equation 22*** and ***Equation 26***) we used a standard fourth-order Runge-Kutta method. At every time step of the numerical integration of the adaptive dynamic equations, the stationary populations of the ecological community (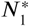 and 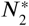) had to be obtained from ***Equation 10***, and their linear stability had to be checked from the Jacobian matrix (***Equation 11***) (***Stucchi et al., 2020***). When a stable stationary population was not found, the integration was aborted; otherwise, the integration was carried on until the evolutionary parameters remained constant for 10^4^ time steps—within a precision of 10^−5^.

## Acknowledgments

This research has been funded by the Spanish Ministerio de Ciencia, Innovación y Universidades-FEDER funds of the European Union support, under projects BASIC (PGC2018-098186-B-I00, J.A.C., and PGC2018-093854-B-I00, J.G. and J.M.P.), and EVA (CGL2016-77377-R, J.M.I).

## Appendix 1

### Adaptive dynamics equations for the two-species ecosystem

According to ***Equation 1***, the per-capita fitness of species *i* is given by

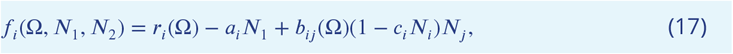

with *r*_*i*_(Ω) and *b*_*ij*_ (Ω) given by ***Equation 6*** and ***Equation 7*** respectively. If the parameters of the mutant are 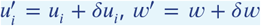, with *δu*_*i*_ = ±1/*K* and *δw* = 0, ±1/*K* (see ***Table 2***), then—given the linearity of *f*_*i*_ with respect to these parameters—the per-capita fitness of the mutant in a steady state community can be written as

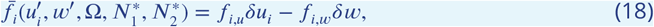

where

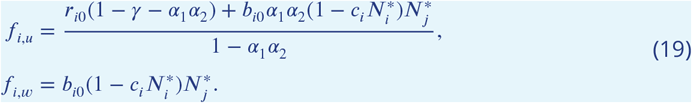

Accordingly, the canonical equations of AD for this sort of mutants read (see ***Dieckmann and Law, 1996***)

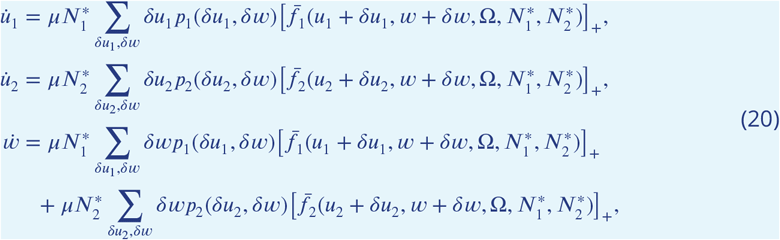

where the sums run over all the corresponding mutations and *p*_*i*_(*δu*_*i*_, *δw*) are their respective probabilities, according to ***Table 2***, and *μ* is the probability of mutation per reproduction event. The function [*x*]_+_ stands for *x* if *x* ≥ 0 and 0 otherwise.

Substituting ***Equation 18*** and ***Equation 19***, performing the sums, and re-scaling evolutionary time with 2*K*^2^/*μ* yields the set of differential equations

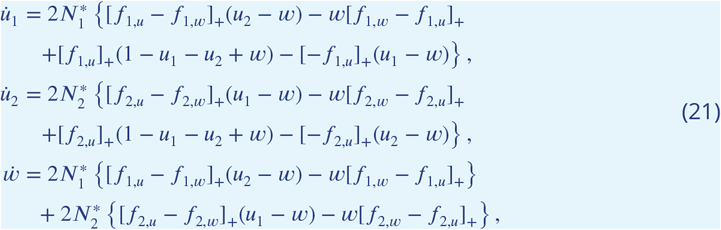

which, using the identity [*x*]_+_ = (*x* + |*x*|)/2, can be rewritten as

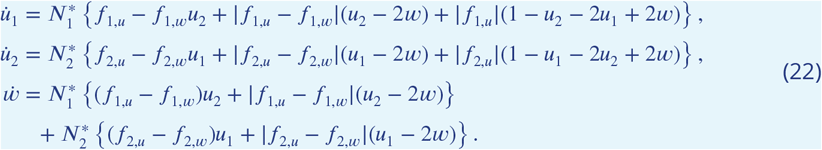

On its side, mutants that change their parameter *α*_*i*_ have a per-capita fitness

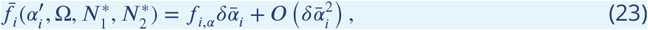

where 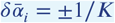 and

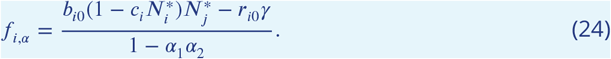

Accordingly,

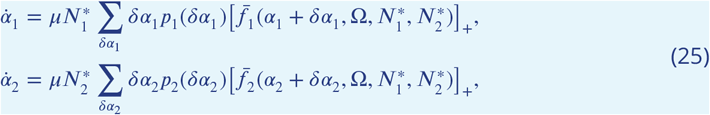

which, neglecting 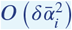 terms and re-scaling time as before, becomes

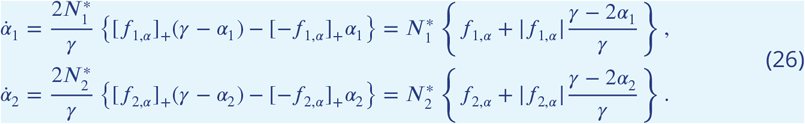

