## Supplementary Material for "Prevalence of mutualism in a simple model of microbial co-evolution"

### Supplementary Information: Prevalence of mutualism in a simple model of microbial co-evolution

#### S1 A toy model for the co-evolution of two microbial species

Consider two microbial species described by the population model

$$\begin{aligned} \dot{X}_1 &= X_1(r_1 - a_1 X_1) + X_1(1 - c_1 X_1) b_{12} X_2, \\ \dot{X}_2 &= X_2(r_2 - a_2 X_2) + X_2(1 - c_2 X_2) b_{21} X_1. \end{aligned} \tag{S1}$$

Let $n_1$ and $n_2$ be the different *exogenous* compounds that they consume, out of a putative set of $K$. A number $q$ of them are common to both species. In their metabolic reactions these microbes yield some byproducts, some of which can in turn be useful for the other species. Let $m_{ij}$ denote the number of compounds endogenously produced by species $j$ that can be used by species $i$ in its own metabolic reactions. Intuitively, the more compounds are used by a microbe the more metabolic reactions can they participate in and the more byproducts ensue. To keep the model simple we will just assume that the number of this products is proportional to that of the compounds involved, irrespective of the species. In other words,

$$m_{21} + \ell_1 = \gamma(n_1 + m_{12}), \qquad m_{12} + \ell_2 = \gamma(n_2 + m_{21}). \tag{S2}$$

However, which fraction correspond to products useful for the other species and which one to useless products is something that is determined by evolution. Thus we assume

$$\begin{aligned} m_{21} &= \alpha_2(n_1 + m_{12}), & \ell_1 &= \frac{\gamma - \alpha_2}{\alpha_2} m_{21}, \\ m_{12} &= \alpha_1(n_2 + m_{21}), & \ell_2 &= \frac{\gamma - \alpha_1}{\alpha_1} m_{12}. \end{aligned} \tag{S3}$$

Solving (S3) leads to

$$m_{21} = \frac{\alpha_2(n_1 + \alpha_1 n_2)}{1 - \alpha_1\alpha_2}, \qquad m_{12} = \frac{\alpha_1(n_2 + \alpha_2 n_1)}{1 - \alpha_1\alpha_2}, \tag{S4}$$

and consequently

$$\ell_1 = \frac{(\gamma - \alpha_2)(n_1 + \alpha_1 n_2)}{1 - \alpha_1\alpha_2}, \qquad \ell_2 = \frac{(\gamma - \alpha_1)(n_2 + \alpha_2 n_1)}{1 - \alpha_1\alpha_2}. \tag{S5}$$

We now need to express the vegetative growth rates $r_1$, $r_2$, as well as the interactions parameters, $b_{12}$, $b_{21}$, in terms of the different number of compounds $n_1$, $n_2$, $m_{21}$, $m_{12}$, $\ell_1$, and $\ell_2$, and the resources shared $q$. For convenience, we will work with fractions, rather than numbers, namely $u_1 = n_1/K$, $u_2 = n_2/K$, $\nu_{21} = m_{21}/K$, $\nu_{12} = m_{12}/K$, and $w = q/K$—the reason being that $K$ will be usually large, and so $\varepsilon = 1/K$ will be a small number.

##### S1.1 Mutations

We now need to model mutations. With a low mutation rate $\mu$, a mutation gets either fixed in the whole population or disappears before a new one shows up. Two kinds of mutations are considered in this model. The first one amounts to changing the number of exogenous compounds that a microbe can use in its metabolic reactions, either adding a new compound to the batch ($n_i \to n_i + 1$, or $u_i \to u_i + \varepsilon$) or eliminating one already present ($n_i \to n_i - 1$, or $u_i \to u_i - \varepsilon$). Any of these two processes may or may not change the resource sharing $q$ by one unit (or $w$ by an amount $\varepsilon$). The probabilities of each possible state transition of the variables $n_1$, $n_2$, and $q$ are gathered in Table 1.

The second kind of mutations will change the number of components that a microbe takes from the other by $\pm 1$. As

$$\frac{\delta\nu_{ij}}{\nu_{ij}} = \frac{\delta\alpha_i}{\alpha_i}, \tag{S6}$$

and $\delta\nu_{ij} = \pm\varepsilon$, we will have

$$\delta\alpha_i = \frac{\pm\varepsilon\alpha_i}{\nu_{ij}}, \tag{S7}$$

| initial state: $n_1, q$ | | | initial state: $n_2, q$ | | |
|---|---|---|---|---|---|
| mutated state | | probability | mutated state | | probability |
| $n_1+1,$ | $q+1$ | $u_2-w$ | $n_2+1,$ | $q+1$ | $u_1-w$ |
| $n_1+1,$ | $q$ | $1-u_1-u_2+w$ | $n_2+1,$ | $q$ | $1-u_1-u_2+w$ |
| $n_1-1,$ | $q$ | $u_1-w$ | $n_2-1,$ | $q$ | $u_2-w$ |
| $n_1-1,$ | $q-1$ | $w$ | $n_2-1,$ | $q-1$ | $w$ |

Table 1: State transitions upon increasing or decreasing one resource from the batch.

so

$$\delta\alpha_1 = \pm\varepsilon\frac{1-\alpha_1\alpha_2}{u_2+\alpha_2 u_1}, \qquad \delta\alpha_2 = \pm\varepsilon\frac{1-\alpha_1\alpha_2}{u_1+\alpha_1 u_2}, \tag{S8}$$

and the probability that $\delta\alpha_i = +\varepsilon$ is $(\gamma-\alpha_i)/\gamma$ and that $\delta\alpha_i = -\varepsilon$ is $\alpha_i/\gamma$.

#### S1.2 Population parameters

We connect the population parameters with in-taken resources and excreted byproducts as follows. As microbes fare better the more resources they have, whereas metabolic byproducts are costly, we assume that the growth rate increases proportional to the amount of consumed resources and decreases with the amount of metabolic byproducts, i.e.

$$\begin{aligned} r_1 &= \frac{r_{10}}{K}(n_1-m_1) = \frac{r_{10}}{1-\alpha_1\alpha_2}\left[(1-\gamma-\alpha_1\alpha_2)u_1-\gamma\alpha_1 u_2\right], \\ r_2 &= \frac{r_{20}}{K}(n_2-m_2) = \frac{r_{20}}{1-\alpha_1\alpha_2}\left[(1-\gamma-\alpha_1\alpha_2)u_2-\gamma\alpha_2 u_1\right]. \end{aligned} \tag{S9}$$

We introduce the factor $K$ for convenience, to express these parameters in terms of the fractions of resources.

Likewise, the interaction coefficients $b_{ij}$ increase with $m_{ij}$, the number of byproducts of species $j$ that are useful to species $i$, and decrease with the competition for the shared resources $q$. Hence,

$$\begin{aligned} b_{12} &= \frac{b_{10}}{K}(m_{12}-q) = b_{10}\left[\frac{\alpha_1}{1-\alpha_1\alpha_2}(\alpha_2 u_1+u_2)-w\right], \\ b_{21} &= \frac{b_{20}}{K}(m_{21}-q) = b_{20}\left[\frac{\alpha_2}{1-\alpha_1\alpha_2}(\alpha_1 u_2+u_1)-w\right]. \end{aligned} \tag{S10}$$

Parameters $r_{i0}$ and $b_{i0}$ are just dimensional constants which set the time scale of the differential equations relative to the mutation rate.

#### S1.3 Adaptive dynamics

According to the population equations (S1), the per capita fitness of both species is given by

$$\begin{aligned} f_1(u_1,u_2,w,\alpha_1,\alpha_2,X_1,X_2) &= r_1(u_1,u_2,\alpha_1,\alpha_2)-a_1X_1+b_{12}(u_1,u_2,w,\alpha_1,\alpha_2)X_2(1-c_1X_1), \\ f_2(u_1,u_2,w,\alpha_1,\alpha_2,X_1,X_2) &= r_2(u_1,u_2,\alpha_1,\alpha_2)-a_2X_2+b_{21}(u_1,u_2,w,\alpha_1,\alpha_2)X_1(1-c_2X_2). \end{aligned} \tag{S11}$$

Thus, at a steady state $X_1=x$, $X_2=y$ (defined by $f_1(u_1,u_2,w,\alpha_1,\alpha_2,x,y)=0$, $f_2(u_1,u_2,w,\alpha_1,\alpha_2,x,y)=0$), the fitness of a mutant will be

$$\bar{f}_i(u_i',w',\alpha_i',u_1,u_2,w,\alpha_1,\alpha_2) = f_{i,u}\delta u_i - f_{i,w}\delta w + f_{i,\alpha}\delta\bar{\alpha}_i, \tag{S12}$$

where

$$\begin{aligned} f_{1,u} &= \frac{r_{01}(1-\gamma-\alpha_1\alpha_2)+b_{01}\alpha_1\alpha_2 y(1-c_1x)}{1-\alpha_1\alpha_2}, \\ f_{1,w} &= b_{01}y(1-c_1x), \\ f_{1,\alpha} &= \frac{b_{01}y(1-c_1x)-r_{01}\gamma}{1-\alpha_1\alpha_2}, \\ f_{2,u} &= \frac{r_{02}(1-\gamma-\alpha_1\alpha_2)+b_{02}\alpha_1\alpha_2 x(1-c_2y)}{1-\alpha_1\alpha_2}, \\ f_{2,w} &= b_{02}x(1-c_2y), \\ f_{2,\alpha} &= \frac{b_{02}x(1-c_2y)-r_{02}\gamma}{1-\alpha_1\alpha_2}, \end{aligned} \tag{S13}$$

and the parameters mutate as $\delta u_i = u_i' - u_i = \pm\varepsilon$, $\delta w = w' - w = \pm\varepsilon$, and $\delta\bar{\alpha}_i = \pm\varepsilon$. Since $\varepsilon \ll 1$, the addition or removal of a compound as a consequence of a mutation has only a marginal effect on the fitness of the mutant, so (S12) makes sense.

The mutant will invade provided $\bar{f}_i > 0$, otherwise it goes extinct. Therefore $\delta u_i$ and $\delta w$, or $\delta\bar{\alpha}_i$, must be so chosen as to make $\bar{f}_i > 0$. Thus, if $f_{i,u} - f_{i,w} > 0$, then $\delta u_i = \delta w = \varepsilon$ is an admissible change, but the opposite change is ruled out. Also if $f_{i,u} > 0$ then mutations like $\delta u_i = \varepsilon$, $\delta w = 0$ are admissible, but the opposite ones are not. Likewise, if the sign of any of those terms is negative, it is the corresponding opposite change that is admissible. A similar argument holds for changes in $\alpha_i$.

Thanks to the linearity of the mutant fitness, the canonical equation of adaptive dynamics for these traits can be readily calculated as

$$\begin{aligned}
\dot{u}_1 &= \mu x\big[\delta u_1 \bar{f}_1(u_1+\delta u_1, w+\delta w, \alpha_1, u_1, u_2, w, \alpha_1, \alpha_2)\big]_+ \\
&= \mu\varepsilon^2 x\left\{[f_{1,u}-f_{1,w}]_+\frac{n_2-q}{K} - \frac{q}{K}[f_{1,w}-f_{1,u}]_+ + [f_{1,u}]_+\frac{K-n_1-n_2+q}{K} - [-f_{1,u}]_+\frac{n_1-q}{K}\right\} \\
\dot{u}_2 &= \mu y\big[\delta u_2 \bar{f}_2(u_2+\delta u_2, w+\delta w, \alpha_2, u_1, u_2, w, \alpha_1, \alpha_2)\big]_+ \\
&= \mu\varepsilon^2 y\left\{[f_{2,u}-f_{2,w}]_+\frac{n_1-q}{K} - \frac{q}{K}[f_{2,w}-f_{2,u}]_+ + [f_{2,u}]_+\frac{K-n_1-n_2+q}{K} - [-f_{2,u}]_+\frac{n_2-q}{K}\right\} \\
\dot{w} &= \mu x\big[\delta w \bar{f}_1(u_1+\delta u_1, w+\delta w, \alpha_1, u_1, u_2, w, \alpha_1, \alpha_2)\big]_+ + \mu y\big[\delta w \bar{f}_2(u_2+\delta u_2, w+\delta w, \alpha_2, u_1, u_2, w, \alpha_1, \alpha_2)\big]_+ \\
&= \mu\varepsilon^2\left\{x\left[[f_{1,u}-f_{1,w}]_+\frac{n_2-q}{K} - \frac{q}{K}[f_{1,w}-f_{1,u}]_+\right] + y\left[[f_{2,u}-f_{2,w}]_+\frac{n_1-q}{K} - \frac{q}{K}[f_{2,w}-f_{2,u}]_+\right]\right\}, \\
\dot{\alpha}_1 &= \mu x\big[\delta\alpha_1 \bar{f}_1(u_1, w, \alpha_1+\delta\alpha_1, u_1, u_2, w, \alpha_1, \alpha_2)\big]_+ = \mu\varepsilon^2 x\left\{[f_{1,\alpha}]_+\frac{\gamma-\alpha_1}{\gamma} - [-f_{1,\alpha}]_+\frac{\alpha_1}{\gamma}\right\}, \\
\dot{\alpha}_2 &= \mu x\big[\delta\alpha_2 \bar{f}_2(u_2, w, \alpha_2+\delta\alpha_2, u_1, u_2, w, \alpha_1, \alpha_2)\big]_+ = \mu\varepsilon^2 y\left\{[f_{2,\alpha}]_+\frac{\gamma-\alpha_2}{\gamma} - [-f_{2,\alpha}]_+\frac{\alpha_2}{\gamma}\right\},
\end{aligned}$$

where $[x]_+ = x$ if $x \geq 0$ and $[x]_+ = 0$ if $x < 0$. And using the fact that $[x]_+ = (x+|x|)/2$ we can rewrite these equations as

$$\begin{aligned}
\dot{u}_1 &= \mu\varepsilon^2 x\left\{(f_{1,u}-f_{1,w})\frac{n_2}{2K} + |f_{1,u}-f_{1,w}|\frac{n_2-2q}{2K} + f_{1,u}\frac{K-n_2}{2K} + |f_{1,u}|\frac{K-n_2-2n_1+2q}{2K}\right\}, \\
\dot{u}_2 &= \mu\varepsilon^2 y\left\{(f_{2,u}-f_{2,w})\frac{n_1}{2K} + |f_{2,u}-f_{2,w}|\frac{n_1-2q}{2K} + f_{2,u}\frac{K-n_1}{2K} + |f_{2,u}|\frac{K-n_1-2n_2+2q}{2K}\right\}, \\
\dot{w} &= \mu\varepsilon^2\left\{x\left[(f_{1,u}-f_{1,w})\frac{n_2}{2K} + |f_{1,u}-f_{1,w}|\frac{n_2-2q}{2K}\right] + y\left[(f_{2,u}-f_{2,w})\frac{n_1}{2K} + |f_{2,u}-f_{2,w}|\frac{n_1-2q}{2K}\right]\right\}, \qquad \text{(S14)} \\
\dot{\alpha}_1 &= \mu\varepsilon^2 x\left\{\frac{1}{2}f_{1,\alpha} + |f_{1,\alpha}|\frac{\gamma-2\alpha_1}{2\gamma}\right\}, \\
\dot{\alpha}_2 &= \mu\varepsilon^2 y\left\{\frac{1}{2}f_{2,\alpha} + |f_{2,\alpha}|\frac{\gamma-2\alpha_2}{2\gamma}\right\}.
\end{aligned}$$

This set of dynamical equations governs the evolution of resources that species 1 and 2 can obtain ($u_1$, $u_2$, respectively), or share ($w$) from the environment, according to their fitness. Note that this evolution occurs in time scale bigger than population dynamics time scale, so that at each integration step a stationary solution of the population dynamics equation (Eq. S1) is found. This new stationary population ($x$, $y$) is used in Eq. S13 to obtain the new fitness $f_{1,2}$ at the next evolution time step.

### S2 Distribution of initial and final interactions

Here we show the distribution of the transitions of species interactions, considering the initial and final states. We present all the cases we explored in Figures 1 to 8. In each figure, we have a set of pie diagrams—one for each value of $\gamma \in \{0.1, 0.2, 0.3, 0.4, 0.5, 0.6, 0.7, 0.8, 0.9\}$—which represent the initial distribution of interactions (left) and the final distribution (right). Figures 1 and 2 correspond to $r_{20} < 1$ and $b_{i0} < 0.01$, figures 3 and 4 correspond to $r_{20} < 0.1$ and $b_{i0} < 0.01$, figures 5 and 6 correspond to $r_{20} < 1$ and $b_{i0} < 0.001$, and figures 7 and 8 correspond to $r_{20} < 0.1$ and $b_{i0} < 0.001$. The species interactions are labelled to the right of each figure to facilitate their identification.

### S3 Stationary populations and evolutionary parameters

Here we show the time evolution of the stationary populations ($N_1^* = x$, $N_2^* = y$) on the right, and their evolutionary parameters on the left, for each one of the examples shown in Figures 6 to 11 of the main article. These parameters

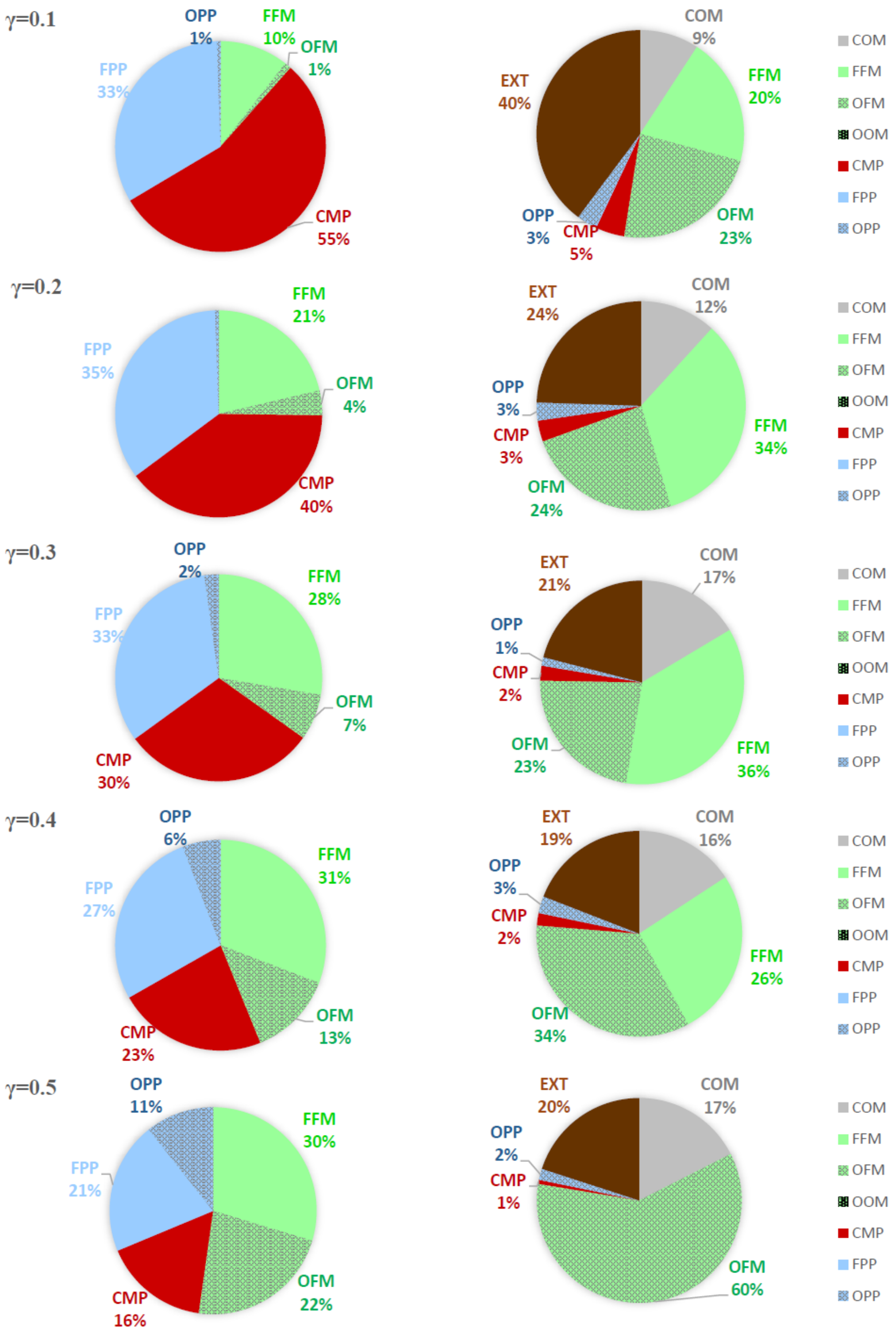


Figure 1: Distribution of initial and final states for parameters: $r_{20} < 1$, $b_{i0} < 0.01$ and $\gamma = 0.1 - 0.5$.

represent the fraction of resources taken by species $i$ ($u_i$), the fraction of shared resources ($w$), and the cross-feeding efficiencies ($\alpha_i$).

Figure 9 shows the transition from competition (CMP) to obligate-facultative mutualism (OFM) including intermediate states of parasitism. The initial parameters were $r_{01} = 1$, $r_{02} = 0.65727$, $b_{01} = 0.00414$, $b_{02} = 0.00447$, $u_1 = 0.17967$, $u_2 = 0.78500$, $w = 0.10050$, $\alpha_1 = 0.08365$, $\alpha_2 = 0.32459$, $\gamma = 0.5$.

Figure 10 shows the transition from facultative parasitism (FPP) to obligate-obligate mutualism (OOM) including intermediate states of mutualism and parasitism. The initial parameters were $r_{01} = 1.0$, $r_{02} = 0.64396$, $b_{01} = 0.00474$, $b_{02} = 0.00262$, $u_1 = 0.60097$, $u_2 = 0.62973$, $w = 0.16897$, $\alpha_1 = 0.13282$, $\alpha_2 = 0.54215$ and $\gamma = 0.6$.

Figure 11 shows the transition from facultative-facultative mutualism (FFM) to commensalism (COM), with inter-

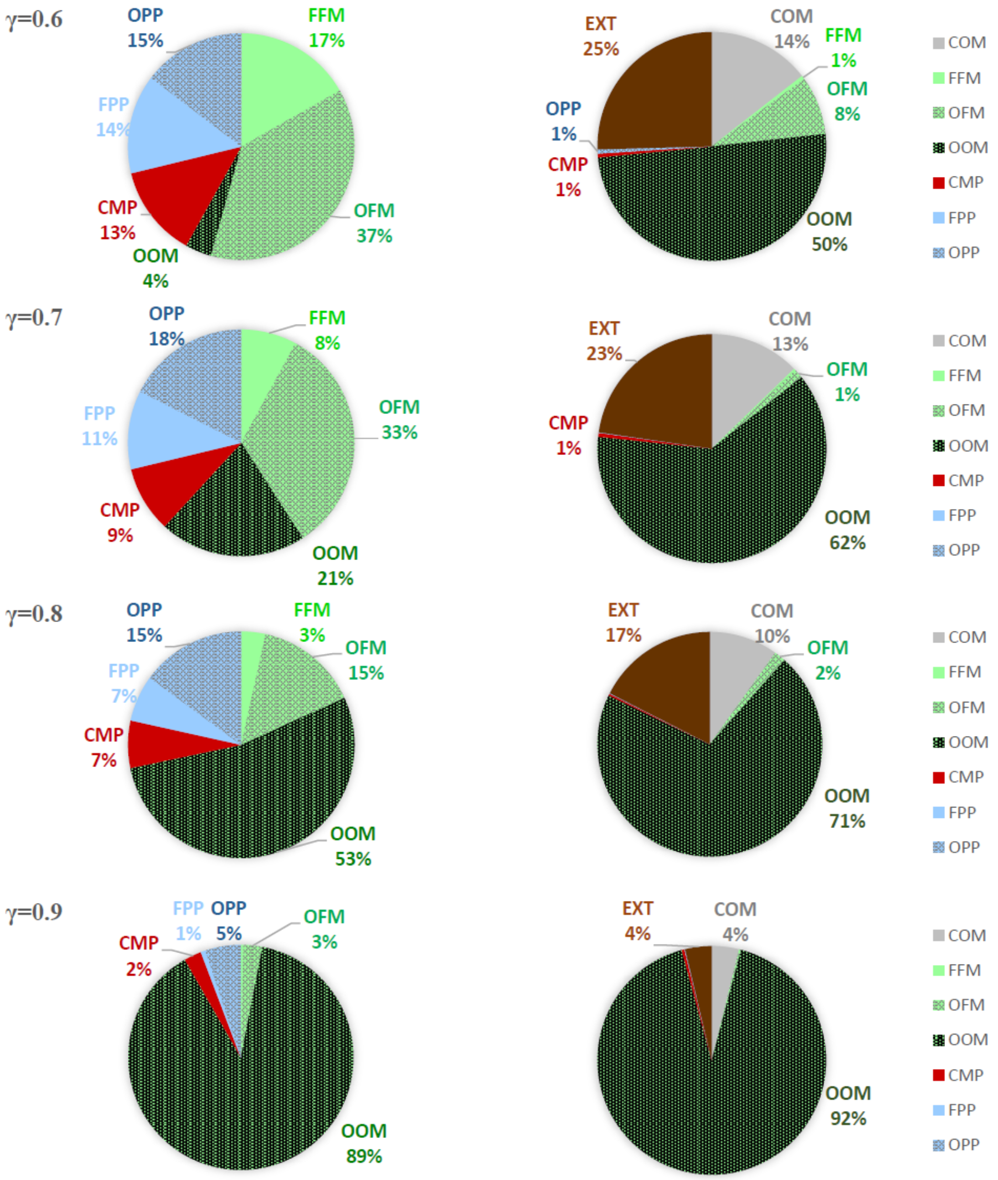


Figure 2: Distribution of initial and final states for parameters: $r_{20} < 1$, $b_{i0} < 0.01$ and $\gamma = 0.6 - 0.9$.

mediate states of parasitism. The initial parameters were $r_{01} = 1.0$, $r_{02} = 0.09827$, $b_{01} = 0.00463$, $b_{02} = 0.00313$, $u_1 = 0.79063$, $u_2 = 0.80318$, $w = 0.00460$, $\alpha_1 = 0.22106$, $\alpha_2 = 0.12717$, and $\gamma = 0.7$.

Figure 12 shows the transition from obligate parasitism (OPP) to obligate-obligate mutualism (OOM), with an intermediate state of mutualism. The initial parameters were $r_{01} = 1.0$, $r_{02} = 0.80306$, $b_{01} = 0.00561$, $b_{02} = 0.00310$, $u_1 = 0.26642$, $u_2 = 0.17242$, $w = 0.01485$, $\alpha_1 = 0.40480$, $\alpha_2 = 0.31492$, and $\gamma = 0.6$.

Figure 13 shows the transition from obligate-facultative mutualism (OFM) to obligate-obligate mutualism (OOM), with intermadiate states of mutualism and parasitism. The initial parameters were $r_{01} = 1.0$, $r_{02} = 0.80306$, $b_{01} = 0.00561$, $b_{02} = 0.00310$, $u_1 = 0.26642$, $u_2 = 0.17242$, $w = 0.01485$, $\alpha_1 = 0.40480$, $\alpha_2 = 0.31492$, and $\gamma = 0.6$.

Figure 14 shows the transition from facultative parasitism (FPP) to obligate-facultative mutualism (OFM), going through an intermediate state of facultative-facultative mutualism (FFM). The initial parameters were $r_{01} = 1.0$, $r_{02} = 1.0$, $b_{01} = 0.00713$, $b_{02} = 0.00309$, $u_1 = 0.38958$, $u_2 = 0.96352$, $w = 0.05325$, $\alpha_1 = 0.10848$, $\alpha_2 = 0.08866$, and $\gamma = 0.2$.

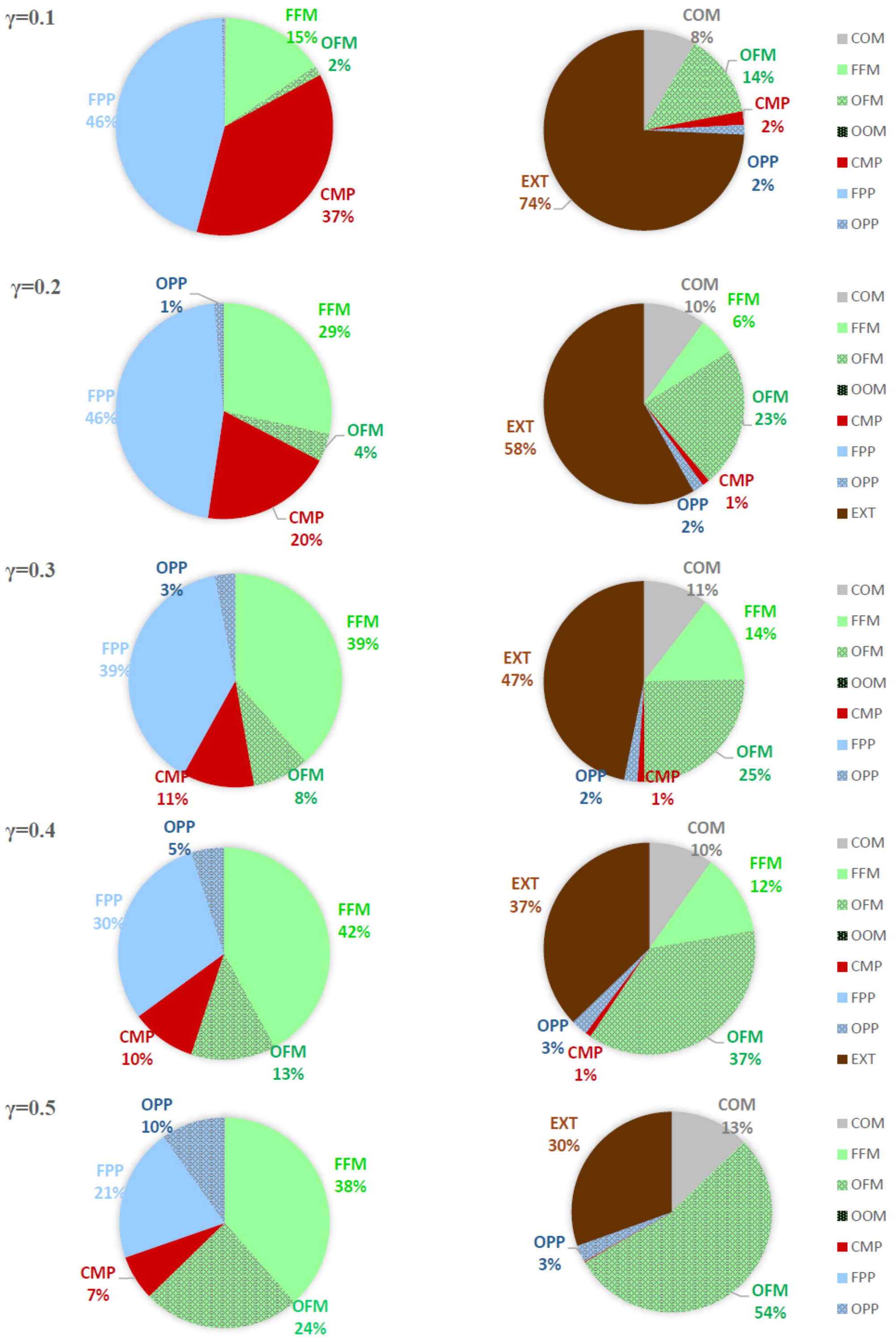


Figure 3: Distribution of initial and final states for parameters: $r_{20} < 0.1$, $b_{i0} < 0.01$ and $\gamma = 0.1 - 0.5$.

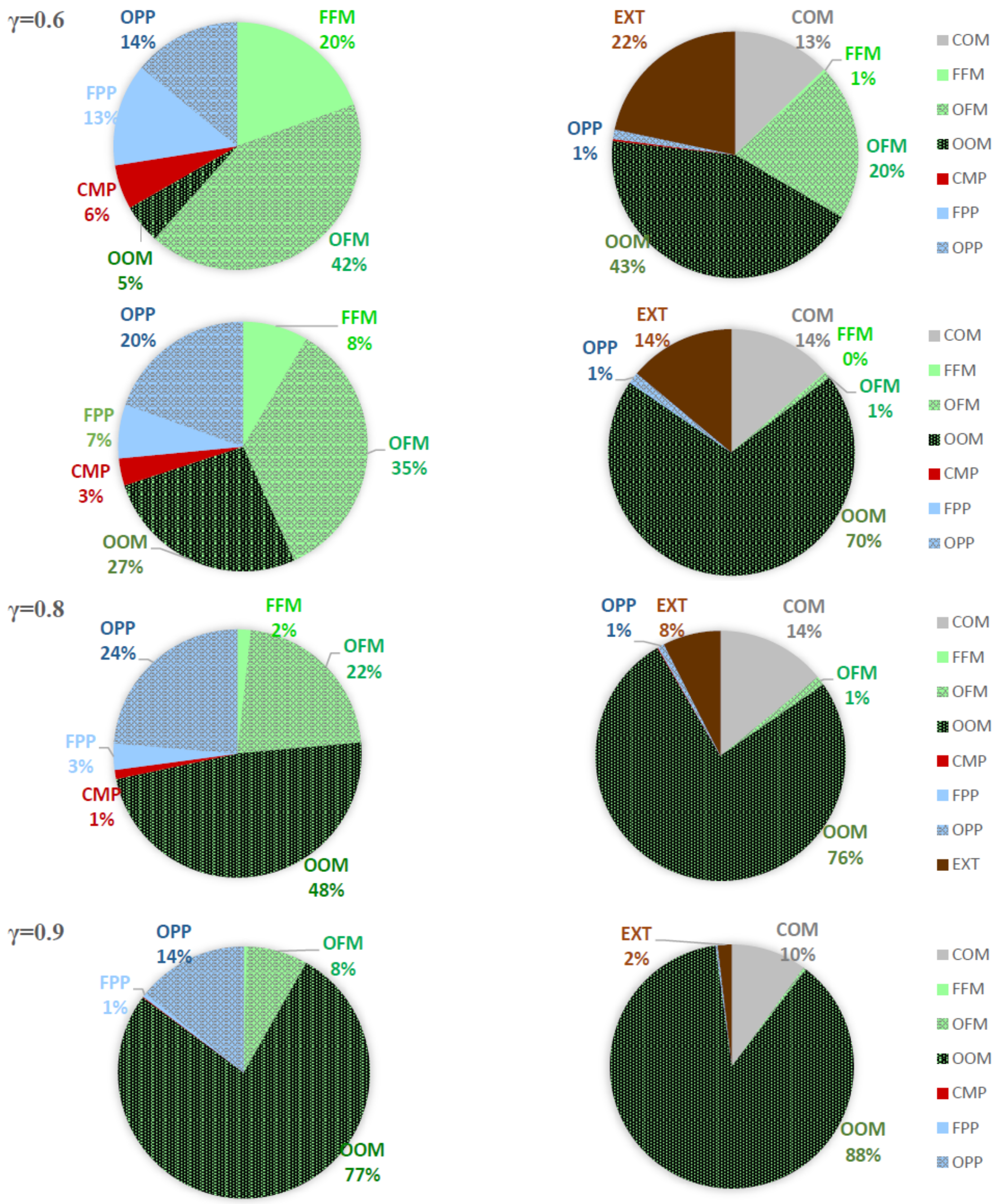


Figure 4: Distribution of initial and final states for parameters: $r_{20} < 0.1$, $b_{i0} < 0.01$ and $\gamma = 0.6 - 0.9$.

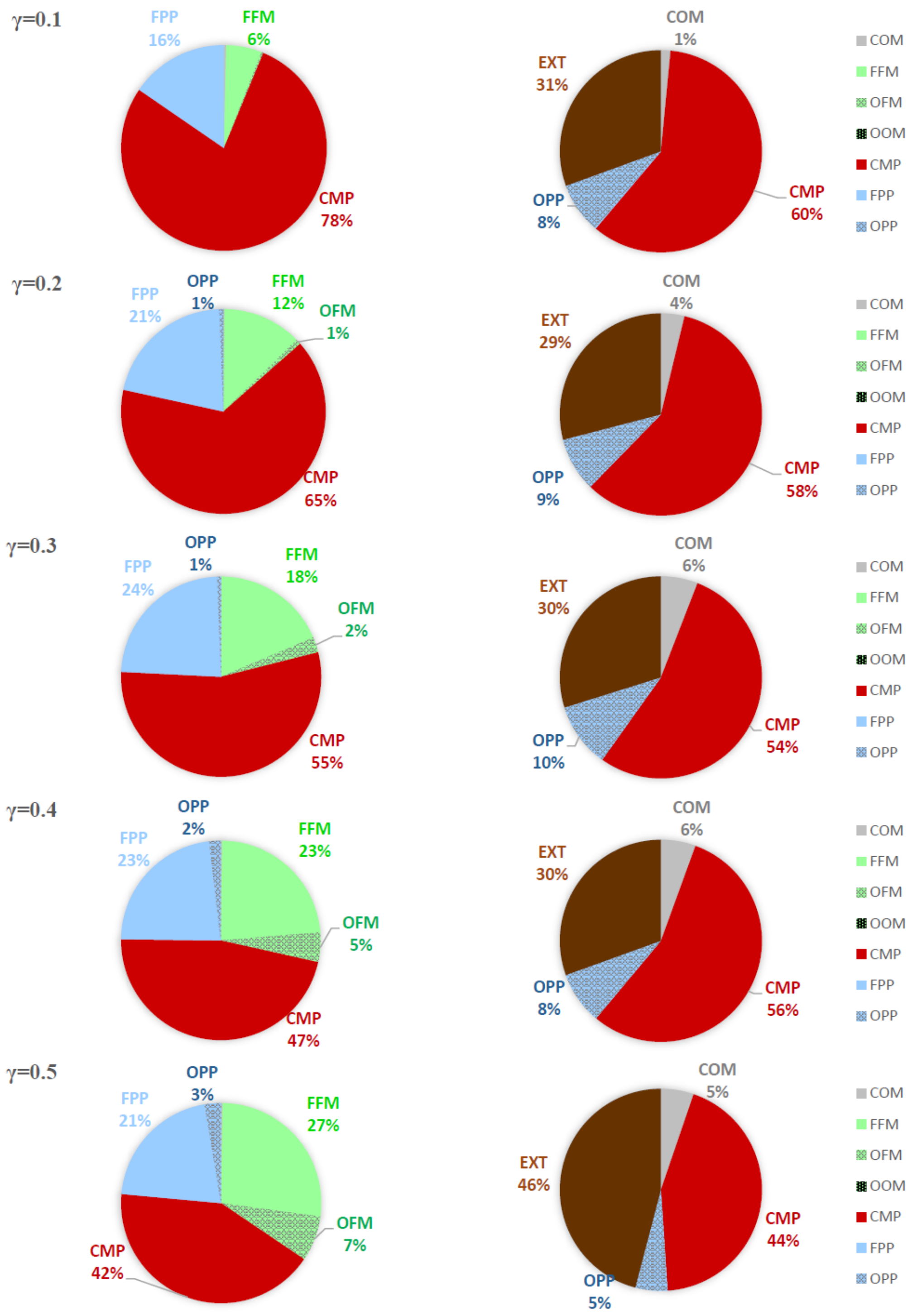


Figure 5: Distribution of initial and final states for parameters: $r_{20} < 1$, $b_{i0} < 0.001$ and $\gamma = 0.1 - 0.5$.

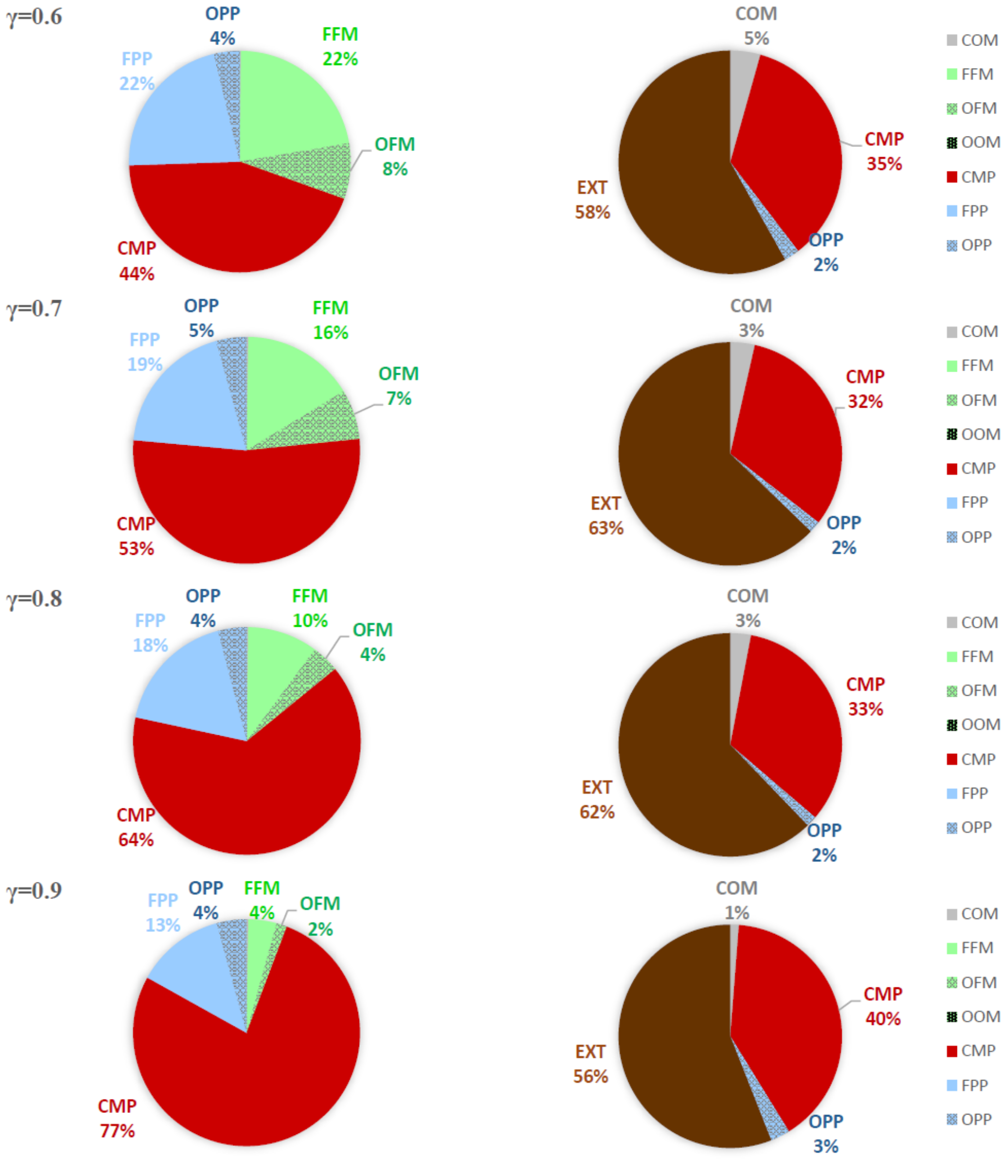


Figure 6: Distribution of initial and final states for parameters: $r_{20} < 1$, $b_{i0} < 0.001$ and $\gamma = 0.6 - 0.9$.

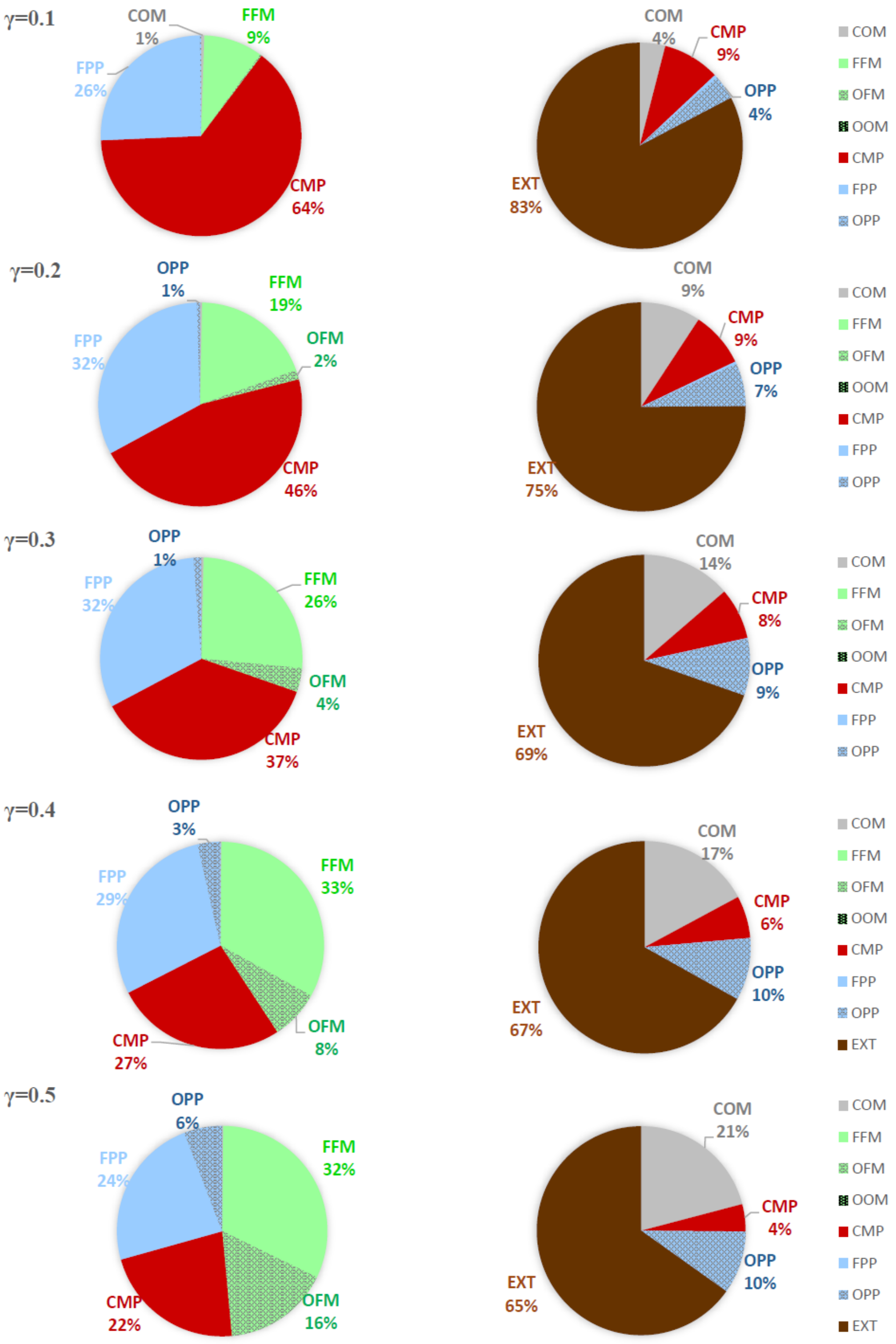


Figure 7: Distribution of initial and final states for parameters: $r_{20} < 0.1$, $b_{i0} < 0.001$ and $\gamma = 0.1 - 0.5$.

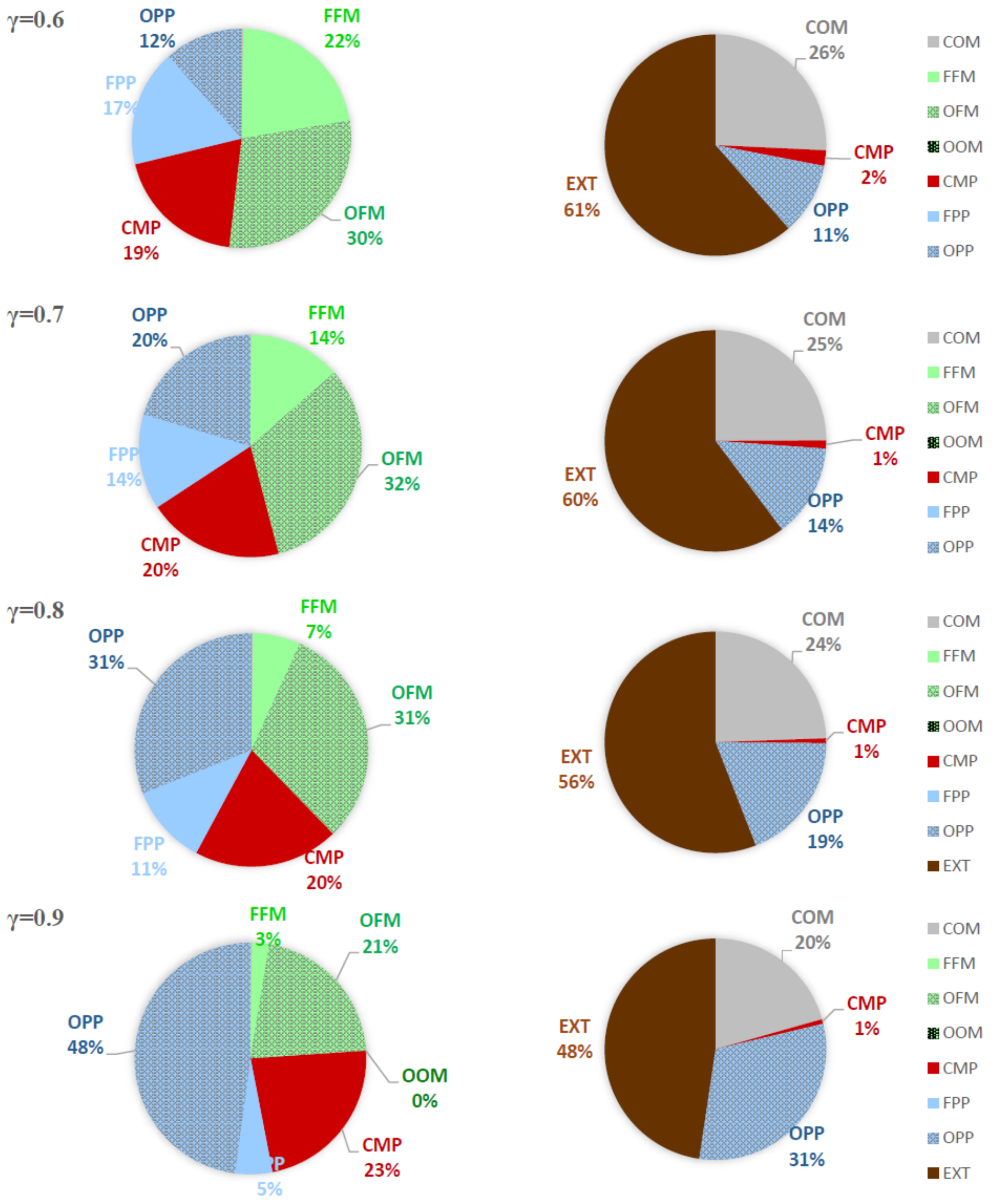


Figure 8: Distribution of initial and final states for parameters: $r_{20} < 0.1$, $b_{i0} < 0.001$ and $\gamma = 0.6 - 0.9$.

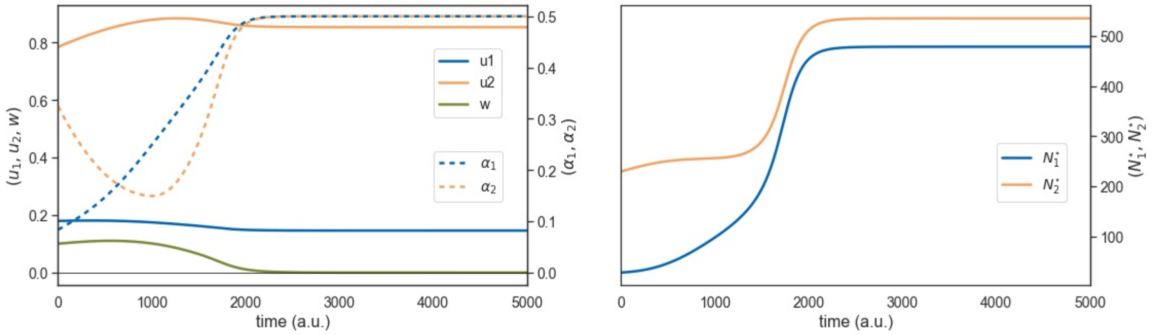


Figure 9: Left: Time evolution of parameters $u_1$, $u_2$, $w$, $\alpha_1$, and $\alpha_2$ for the co-evolution shown in Figure 6. Right: Stationary population along the evolution.

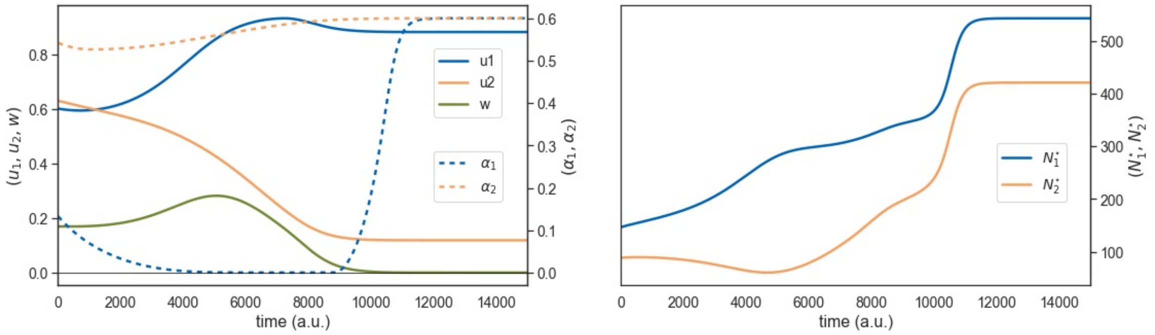


Figure 10: Left: Time evolution of parameters $u_1$, $u_2$, $w$, $\alpha_1$, and $\alpha_2$ for the co-evolution shown in Figure 7. Right: Stationary population along the evolution.

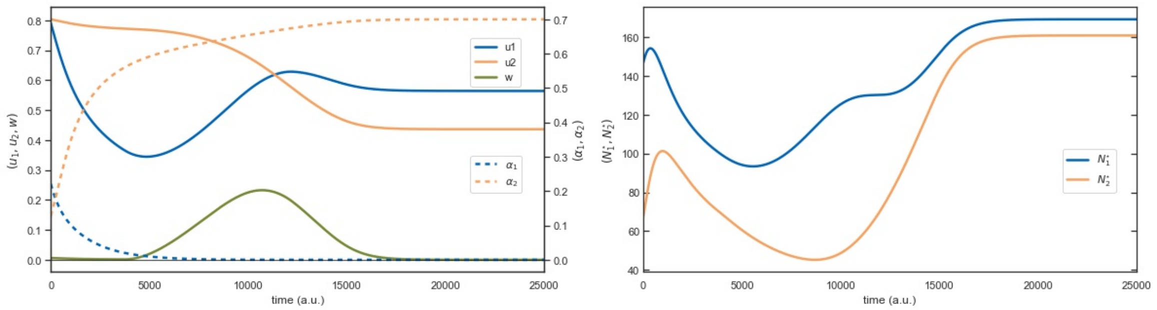


Figure 11: Left: Time evolution of parameters $u_1$, $u_2$, $w$, $\alpha_1$, and $\alpha_2$ for the co-evolution shown in Figure 8. Right: Stationary population along the evolution.

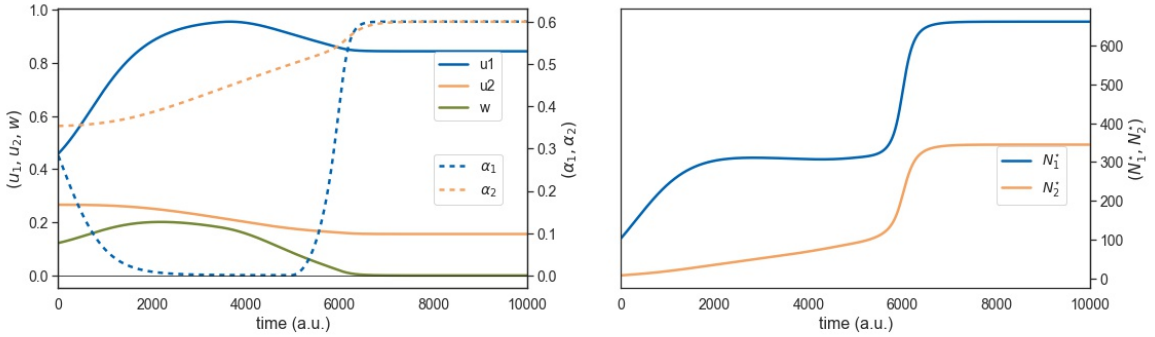


Figure 12: Left: Time evolution of parameters $u_1$, $u_2$, $w$, $\alpha_1$, and $\alpha_2$ for the co-evolution shown in Figure 9. Right: Stationary population along the evolution.

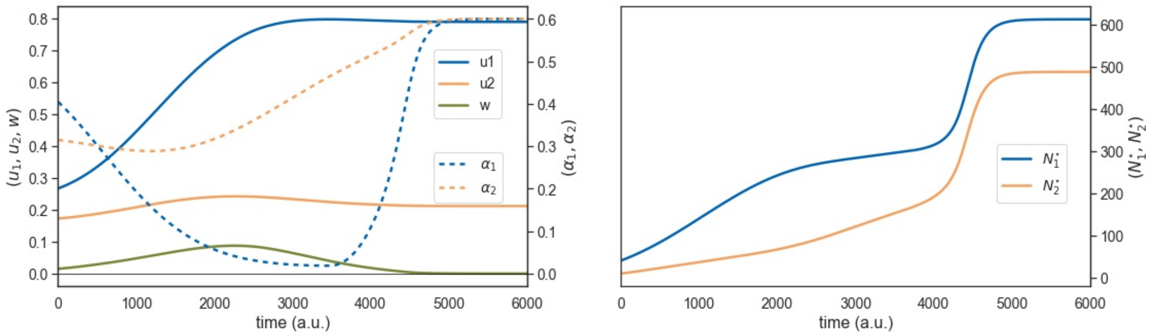


Figure 13: Left: Time evolution of parameters $u_1$, $u_2$, $w$, $\alpha_1$, and $\alpha_2$ for the co-evolution shown in Figure 10. Right: Stationary population along the evolution.

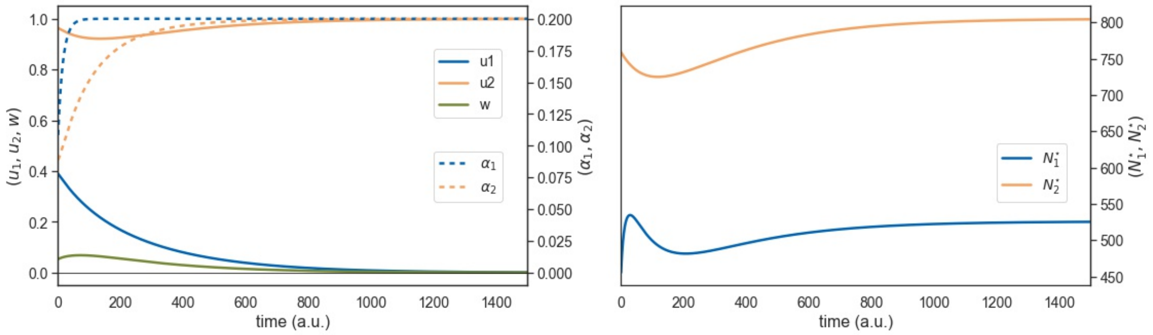


Figure 14: Left: Time evolution of parameters $u_1$, $u_2$, $w$, $\alpha_1$, and $\alpha_2$ for the co-evolution shown in Figure 11. Right: Stationary population along the evolution.
